# Set2 and H3K36 regulate the *Drosophila* male X chromosome in a context-specific manner, independent from MSL complex spreading

**DOI:** 10.1101/2024.05.03.592390

**Authors:** Harmony R. Salzler, Vasudha Vandadi, A. Gregory Matera

## Abstract

Dosage compensation in *Drosophila* involves upregulating male X-genes two-fold. This process is carried out by the MSL (male-specific lethal) complex, which binds high-affinity sites and spreads to surrounding genes. Current models of MSL spreading focus on interactions of MSL3 (male-specific lethal 3) with histone marks; in particular, Set2- dependent H3 lysine-36 trimethylation (H3K36me3). However, Set2 might affect DC via another target, or there could be redundancy between canonical H3.2 and variant H3.3 histones. Further, it is difficult to parse male-specific effects from those that are simply X-specific. To discriminate among these possibilities, we employed genomic approaches in *H3K36* (residue) and *Set2* (writer) mutants. The results confirm a role for Set2 in X-gene regulation, but show that expression trends in males are often mirrored in females. Instead of global male-specific reduction of X-genes in *Set2/H3K36* mutants, the effects were heterogeneous. We identified cohorts of genes whose expression was significantly altered following loss of H3K36 or Set2, but the changes were in opposite directions, suggesting that H3K36me states have reciprocal functions. In contrast to *H4^K16R^* controls, analysis of combined *H3.2^K36R^/H3.3^K36R^* mutants neither showed consistent reduction in X-gene expression, nor any correlation with MSL3 binding. Examination of other developmental stages/tissues revealed additional layers of context-dependence. Our studies implicate BEAF-32 and other insulator proteins in Set2/H3K36-dependent regulation. Overall, the data are inconsistent with the prevailing model wherein H3K36me3 directly recruits the MSL complex. We propose that Set2 and H3K36 support DC indirectly, via processes that are utilized by MSL but common to both sexes.

## Introduction

The evolution of heterogametic sexes necessitates that the number of X chromosome transcripts from XY males and XX females be equalized to prevent maladaptive disparities in gene dosage. In mammals, this dosage compensation (DC) system involves stochastic inactivation of one female X chromosome [1]. In contrast, *Drosophila melanogaster* relies on a roughly 2-fold upregulation of transcripts generated from the male X. Importantly, many elements of the *Drosophila* DC system are conserved in mammals [2], and relevant to human health and disease research [3–5].

The most extensively studied mediator of DC in *Drosophila* is the Male-Specific Lethal (MSL) complex, which carries out histone H4 lysine 16 acetylation (H4K16ac), primarily on the male X [6, 7]. One estimate suggests that the MSL complex accounts for ∼40-50% of the upregulation of the male X [8]. Genetic mutations in MSL complex members demonstrate that it is essential for male survival [9–11]. Current evidence supports involvement of the MSL complex in regulating RNA polymerase II elongation [12–14] as well as in genome organization [14–18]. Importantly, recent work also demonstrates that the H4K16 residue itself is essential in male flies, and that the H4K16 acetylation function of the MSL complex is crucial [19, 20].

The core MSL complex is comprised of five proteins (MSL1, MSL2, MSL3, MLE, and MOF) and two lncRNAs (*roX1* and *roX2)* [14, 21]. Four of the five MSL proteins are also present in females, excepting MSL2 [22, 23]. The MOF acetyltransferase, which catalyzes acetylation of H4K16ac, also acts on housekeeping genes throughout the genome in the context of the non-specific lethal (NSL) complex [24]. The distributions of H4K16ac resulting from these two complexes are distinct, as MSL acetylates over gene bodies, whereas NSL preferentially targets promoters [25, 26]. Other MSL-interacting proteins have been identified, [27], many of which have substantiated roles in DC [27–30].

Current models of MSL function posit that the complex is initially targeted to the male X via binding of MSL2•MSL1 dimers to high-affinity binding sites (HASs), followed by subsequent spreading to nearby genes [31](for reviews see [14, 21]). The CLAMP protein is an important cofactor for MSL2•MSL1 binding [32, 33], although CLAMP-independent binding to a small subset of so-called PionX (pioneering on the X) sites is required for initial recognition of the male X [34]. Following initial targeting, MSL activity spreads to surrounding active genes by way of the MSL3 chromodomain [35, 36].

To date, our understanding of how MSL3 facilitates spreading to nearby active genes remains incomplete and controversial. Early evidence pointed to the importance of histone H3 lysine-36 trimethylation (H3K36me3) and its cognate lysine methyltransferase, Set2, in propagating the MSL complex across the male X. First, *Set2* null male larvae exhibit a 2-10 fold reduction in MSL complex recruitment to a subset of X-genes [37]. Second, recombinant MSL3 displays an affinity for H3K36me3 modified nucleosomes [37]. Despite these findings, MSL recruitment defects observed in *Set2* mutants were inconsistent regarding H4K16ac and/or mRNA levels over the genes examined [37]. Furthermore, a plasmid model of DC also called into question the importance of H3K36me3 [38].

More recently, RNA-seq analysis of *Set2* mutant male larvae substantiated a small, but significant decrease in X-gene expression, but the same study also found that *H3.2^K36R^* and *H3.3B;H3.3A* null mutants failed to display this effect [39]. Given that many histone methyltransferases are known to target non-histone substrates [40–42], including the mammalian ortholog of Set2 (SETD2) [43–45], it is plausible that the effect of *Set2* loss on male X-expression is mediated by a target other than H3K36. However, other plausible interpretations of these data remain.

The absence of females in previous studies also makes it difficult to discern whether global X chromosome effects in male cells are due to “maleness” or “X-ness” in the sense that the X itself has unique features not specific to sex that could impact gene regulation [46–50]. Furthermore, the issue of functional redundancy between H3.2 and H3.3 K36 residues [51] was not considered [39]. Finally, with respect to DC, the potential for heterogeneous regulation of X-genes has been underexplored. In particular, work investigating “non-canonicial” DC mechanisms provide important hints that mechanisms for balancing sex chromosome gene dosage may not be entirely mediated by the MSL complex [8, 18].

In this study, we utilize histone genetics and transcriptome profiling to clarify the relationship between Set2, H3K36me3, MSL3 recruitment, and X chromosome gene regulation. We confirm previous reports that Set2 impairs gene expression on the X chromosome. However, our inclusion of females combined with nuanced bioinformatic analyses reveal that the effects of Set2, H3.2K36, and H3.3K36 on X chromosome gene expression are surprisingly heterogeneous. Importantly, our analysis of an *H3.3^K36R^/H3.2^K36R^* combined mutant addresses the possibility of functional redundancy between histone variants, and we find no evidence of involvement of H3K36 on promoting expression of dosage-compensated genes. Interestingly, we frequently observe opposite effects on gene expression between *Set2* and *H3^K36R^* mutants at multiple developmental stages suggestive of a regulatory switch between methyl states of H3K36. Lastly, we find that X-genes with decreased expression in *Set2* and *H3^K36R^* mutants in larval brain are enriched in components of the BEAF-32 insulator complex compared to unaffected genes.

Based on these findings, we conclude that neither *Set2* nor *H3K36* are required for MSL3 recruitment, as their effects are gene-specific, context-dependent, and do not reliably correlate with the presence MSL3 binding or H4K16ac. Rather, we argue that the evidence is more compatible with *Set2* mediated H3K36 trimethylation impacting other processes utilized in DC, but not specific to DC (such as elongation control or 3D genome organization).

## Results

### *H3.2^K36R^* and *H3.3^K36R^* mutations do not specifically impair male viability

Male-specific lethality is a defining feature of mutations that affect DC in *Drosophila* (reviewed in [21, 52]). Remarkably, this specificity extends all the way down to the histone residues themselves, as an *H4^K16R^* mutation causes developmental delay and death in male progeny whereas their female siblings are completely viable [19]. This male-specific lethality can be bypassed by expression of an acetylation mimicking *H4^K16Q^* mutation [20]. Together, these results demonstrate that H4K16ac is the critical PTM of the DC machinery in *Drosophila*. Moreover, they show that H4K16 is not required for basal genome function, as female gene expression and viability were unaffected.

If H3K36me3 plays an important role in the localization or spreading of the MSL complex, one might expect to observe decreased male viability in mutants that inhibit H3K36 methylation. To test this idea, we assayed the fraction of adult males in H3.2 and H3.3 K36R mutants, along with H4 K16R and HWT (histone wild type) controls. For complete genotypes and genetic schemes for generating these animals see Figures 1A, S1 and Table S1. Note that *Set2* null and 12x*H3.2^K36R^* animals fail to eclose as adults, but wandering L3 males from these lines are readily obtained [37, 53]. To ascertain whether H3.2 K36 and H4 K16 residues interact genetically, we carried out complementation analysis between multi-gene families [51]. That is, we combined two 12x histone constructs *in trans,* and assayed pupation and eclosion frequencies of the resulting progeny. A significant change in viability by comparison to control crosses would suggest that the two residues cooperate in common pathways. Previously, we found that H3.2 K36R interacted strongly with K27R but was fully complemented by a K9R mutation [51].

**Fig. 1.**
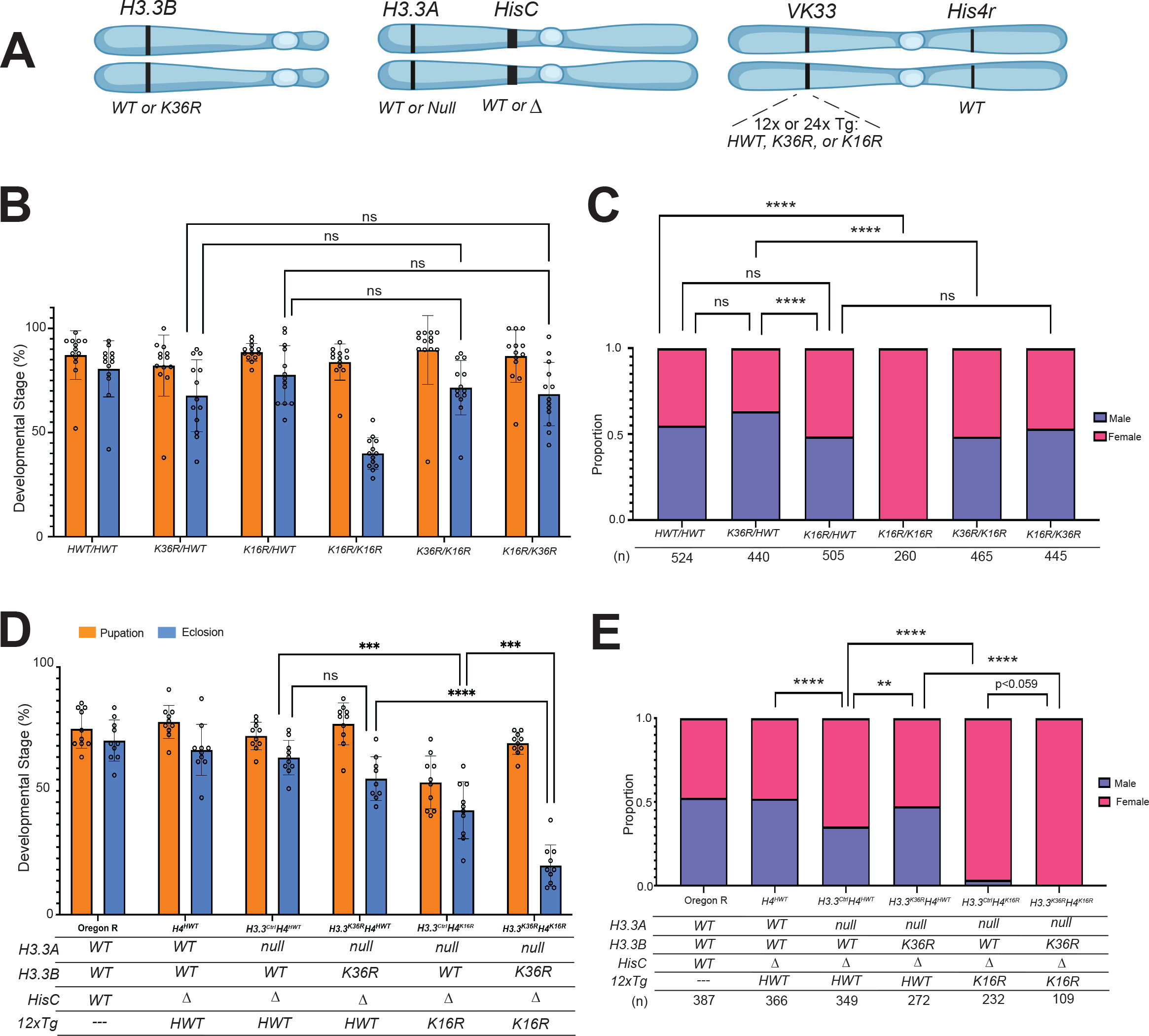
H**3.3K36R interacts genetically with H4K16R. (**For all genotypes in B&C, the *H3.3A* gene (chr. 2L) and *H3.3B* gene (chr. X) are WT, and the endogenous replication-dependent histone gene cluster, *HisC* (chr. 2L) is Δ. The transgenic insertion site *VK33* (chr. 3L, band 65B2) was used for all 12x histone transgenes. For each genotype, 2 copies of *12x* transgenes (*HWT, H3.2K36R,* or *H4K16R*) are present in trans. All C&D genotypes, except Oregon R (OR), are *HisC*Δ. Status of *H3.3A, H3.3B,* and *12x* transgenes are indicated in the table below the graph in (D). *H3.3B* is either WT or K36R; *H3.3A* WT or null. *HisC* is either intact (WT) or Δ (null). *12xH4* transgenes contain 12 copies of the histone repeat unit, each containing all five replication-dependent histone genes. Transgenes used in this study carry the following alleles of *H4*: *HWT* or *K16R*. Cartoon of genetic loci used in panel D. Panels A and C were created using BioRender.com. **A)** Cartoon of genetic loci used in panels B-D. For complete genotypes, see Figs. S1, S2, S4, & S5. **(B)** Developmental viability assay. For each genotype, % pupation and % eclosion of 8-10 biological replicates (50 larvae/replicate vial) were calculated, and means and SD of these percentages were plotted. Statistical significance for % eclosion was calculated with GraphPad Prism software using Brown-Forsythe and Welch ANOVA tests, followed by the Dunnett’s T3 multiple comparisons test. \*\*\**P* < 0.001. \*\*\*\**P* < 0.0001. ns, not significant. **(C)** Proportion of male and female eclosed animals were calculated. Statistical significance for sex ratio was calculated with GraphPad Prism software using Fisher’s Exact Test, followed by the Benjamini-Hochberg False Discovery Rate (FDR) correction for multiple comparisons (Q=0.05). \*\**P* < 0.01. \*\*\*\**P* < 0.0001. ns, not significant. **(D)** Viability assay, as in B. **(E)** Sex ratio of adults, as in C.

As shown in Fig. 1B, addition of an *HWT* transgene fully rescued the larval and pupal viability defects seen in *K36R*-only animals [51]. However, there was no significant change in the number of males that eclose from a *K36R/HWT* cross compared to *HWT/HWT* controls (Fig. 1C). If anything, there was a modest *increase* in *K36R/HWT* adult males. Consistent with its known role in DC, we did observe a slight but insignificant decrease in the fraction of males emerging from a *K16R/HWT* cross compared to the control (Fig. 1C). However, the opposing sex skew of the *K36R/HWT* and *K16R/HWT* adults resulted in a statistically significant difference (Fig. 1C). We also observed that modifying the *K36R/HWT* genotype to *K36R/K16R* resulted in a significant decrease in males, but the converse was not true (Fig. 1C). Modification of *K16R/HWT* to *K16R/K36R* resulted in no change (Fig. 1C). Together, these two observations imply that the male-diminishing effect of *K16R* predominates over the male-promoting effect of *K36R*. Thus H3.2K36R histones appear to be slightly more toxic to females, whereas H4K16R histones specifically affect males.

Importantly, we note the significant absence of adult males in *K16R/K16R* crosses, despite the presence of wildtype copies of *His4r* in this background (Fig. 1C). Although *His4r* is a replication-independent histone gene, it expresses an identical H4 protein. Previously, we found that animals bearing a single 12x *K16R* transgene (crossed in maternally) in a *His4r* positive background resulted in 8.5% eclosed males [19]. Taken together, these findings support the notion that the proportion of zygotically expressed H4, compared to the amount of wild-type maternal histones and His4r, is a critical determinant of male viability.

In contrast with the results for H3.2, we found that *H3.3^K36R^* mutants complete development and eclose at a frequency of ∼80%, which is nearly identical to that of *H3.3^Ctrl^* animals [51] (for full genotypes see Fig S2). We therefore assessed the ratio of males and females in adults of these genotypes. We found that *H3.3^K36R^* males comprise ∼50%, of eclosed adults, which is slightly but not significantly greater than that of the *H3.3^Ctrl^* (Fig. S3). These data suggest that an *H3.3^K36R^* mutation does not substantially weaken dosage compensation.

### *H3.3^K36R^* interacts genetically with *H4^K16R^*

Synthetic lethal (or synthetic sick) interactions are those wherein the combination of two different mutations produces death or other strong phenotypes, whereas single mutations do not. Synthetic interactions can thus implicate two genes as participating in a common pathway [54, 55]. Given the importance of H4K16ac to *Drosophila* DC, we wondered whether genetic evidence for involvement of H3.3K36 in DC might emerge in the sensitized background of an *H4^K16R^* mutation.

We hypothesized that if H3.3K36 were involved in DC, the male lethal phenotype of the *H4^K16R^* mutant would be enhanced. We therefore assayed overall viability and male:female ratios in genotypes combining *H3.3^Ctrl^* and *H3.3^K36R^* mutations with *H4^K16R^* (Fig 1A) (For full genotypes, see Fig. 4). In these experiments, *His4r* was wild type, as deletion of this locus rendered the *H4^K16R^* male lethal phenotype too severe to detect synthetic effects (32% adult males vs 0%; see [19]). As expected, overall viability levels for Oregon R (*OreR*), *H4^HWT^*, and *H3.3^Ctrl^H4^HWT^* control genotypes were similar for both pupation and eclosion (Fig. 1D). The addition of *H3.3B^K36R^* to generate *H3.3^K36R^H4^HWT^* animals had no significant impact on viability, though recent work shows that this mutation does reduce adult lifespan (Fig. 1D, [56]). In contrast, *H3.3^Ctrl^H4^K16R^* mutants exhibited a significant reduction in viability (∼45% eclosion). This value is comparable to the eclosion frequency reported for *H4^K16R^* animals bearing wild type *H3.3* genes (50%) (Fig. 1D, [19]). Interestingly, when *H3.3^K36R^* and *H4^K16R^* mutations are combined, adult survival is severely impaired (∼20%; see Fig. 1D), strongly suggesting that H3.3K36 and H4K16 regulate common pathways. However, the degree of synthetic lethality also suggests that both males and females are affected.

Given that H4K16ac is also deposited in the context of autosomal promoters, we examined whether there was a more severe viability defect in males, suggestive of an impairment to DC. We calculated the proportions of males and females from the eclosed viable adults. As expected, *OreR* and *H4^HWT^* produced roughly equal numbers of males and females, but the *H3.3^Ctrl^H4^HWT^* control skewed significantly female (Fig. 1E). We note that this imbalance was unexpectedly ‘rescued’ by mutation of *H3.3B^K36R^* (*H3.3^K36R^H4^HWT^*; Fig. 1E), suggesting that loss of H3.3K36 can promote male survival in the context of H3.3 insufficiency. Strikingly, the *H3.3^Ctrl^H4^K16R^* genotype exhibited dramatic impairment of male survival, despite the presence of a wild-type *His4r* gene. Compared to previous reports, ablation of *H3.3A* reduced male survival 10-fold in the context of an *H4^K16R^* mutation (3.4%, Fig. 1E compared to 32%, [19]). Interestingly, combining *H3.3^K36R^* and *H4^K16R^* mutations (*H3.3^K36R^H4^K16R^*) completely eliminated eclosion of viable males. This finding is consistent with the possibility that H3.3K36 performs a role in DC, however, given that females were also affected to a lesser extent, the possibility that combining these mutations confers a global reduction in viability that disproportionately affects weakened males cannot be excluded.

### Transcriptomic analysis of *Set2* and *H3^K36^* mutants in the larval brain

Although the genetic interaction between *H3.3^K36R^* and *H4^K16R^* was intriguing, we wanted to assay the effects of K36 residue and writer mutations on male and female transcriptomes. A previous study had analyzed brains of male *Set2^1^* (a null allele), *H3.3^WT^H3.2^K36R^* and *H3.3Δ* (*H3.3B^null^;H3.3A^null^*) wandering 3^rd^ instar (WL3) larvae [39]. These investigators identified a role for Set2 in supporting expression of X chromosome transcripts in males, however the exclusion of females from that study makes it unclear if this effect is truly male-specific or simply X-specific. Moreover, the complete absence of H3.3 protein removes an important nucleosomal subunit from many different subcompartments of the genome, presumably replacing it with wildtype H3.2.

To extend the analysis to females and to better parse the relative involvement of Set2, H3.2K36, and H3.3K36 in the regulation of gene expression, we performed poly-A selected RNA-seq followed by DESeq2 differential expression analyses in WL3 brains. Altogether, there were six replicates (3 male and 3 female) of three different mutant genotypes plus three corresponding controls: *Set2^1^* and *yw*; *H3.3^WT^H3.2^K36R^* and *H3.3^WT^H3.2^HWT^*; *H3.3^K36R^H3.2^HWT^* and *H3.3^Ctrl^H3.2^HWT^* (see Fig. S5 and Table 1 for detailed descriptions). Note that we analyzed the *H3.3B^K36R^* mutation on the *H3.2^HWT^* histone replacement background to enable direct comparison with the *H3.3^WT^H3.2^K36R^* animals. We also sequenced samples to high read depth (62-95 million paired-end reads per replicate) and avoided cutoffs based on a log2 fold-change (LFC) thresholds in downstream analyses because previous work has shown that mutation and knockdown of MSL complex members yield subtle LFC values X chromosome-wide [31, 57].

Principal component analysis (PCA) revealed tight groupings of replicates by genotype, as well as by sex (Fig. S6A). For our initial DESeq2 runs, we combined replicates for both sexes into a single genotype class to simplify general trends in expression patterns between the mutants. MA plots highlighting all differentially expressed genes (DEGs) (adjusted *P* value < 0.05) revealed a notably greater number of DEGs in the *Set2^1^* mutant (7,042) than either the *H3.3^WT^H3.2^K36R^* (4,519) or the *H3.3^K36R^H3.2^HWT^* (1,835) mutant alone, or their sum (6,344) (Fig. 2A). When adjusting this sum to account for genes that are DEGs in both H3K36 mutant genotypes (5,508 for one or both H3K36R mutants; Fig S6B), these data not only suggest significant functional compensation between H3.2K36 and H3.3K36, but also the possibility of Set2 functions that are not related to H3K36. This pattern was maintained when an LFC cutoff of > |1| was employed (Fig. S6C). We also note that, within the subset of DEGs identified in all three mutant genotypes (618 genes), the largest group of genes was upregulated in all three mutants (43%, see Fig. S6B). Additionally, a substantial fraction (25%) was upregulated in both H3K36R mutant genotypes, but downregulated in the *Set2^1^* mutant, suggesting a regulatory relationship between H3K36 trimethylation and other modification states (Fig. S6B). Importantly, these data hint at other possible regulatory scenarios besides H3K36-independent functions of Set2 or redundancy between H3.2 and H3.3 residues.

**Fig. 2.**
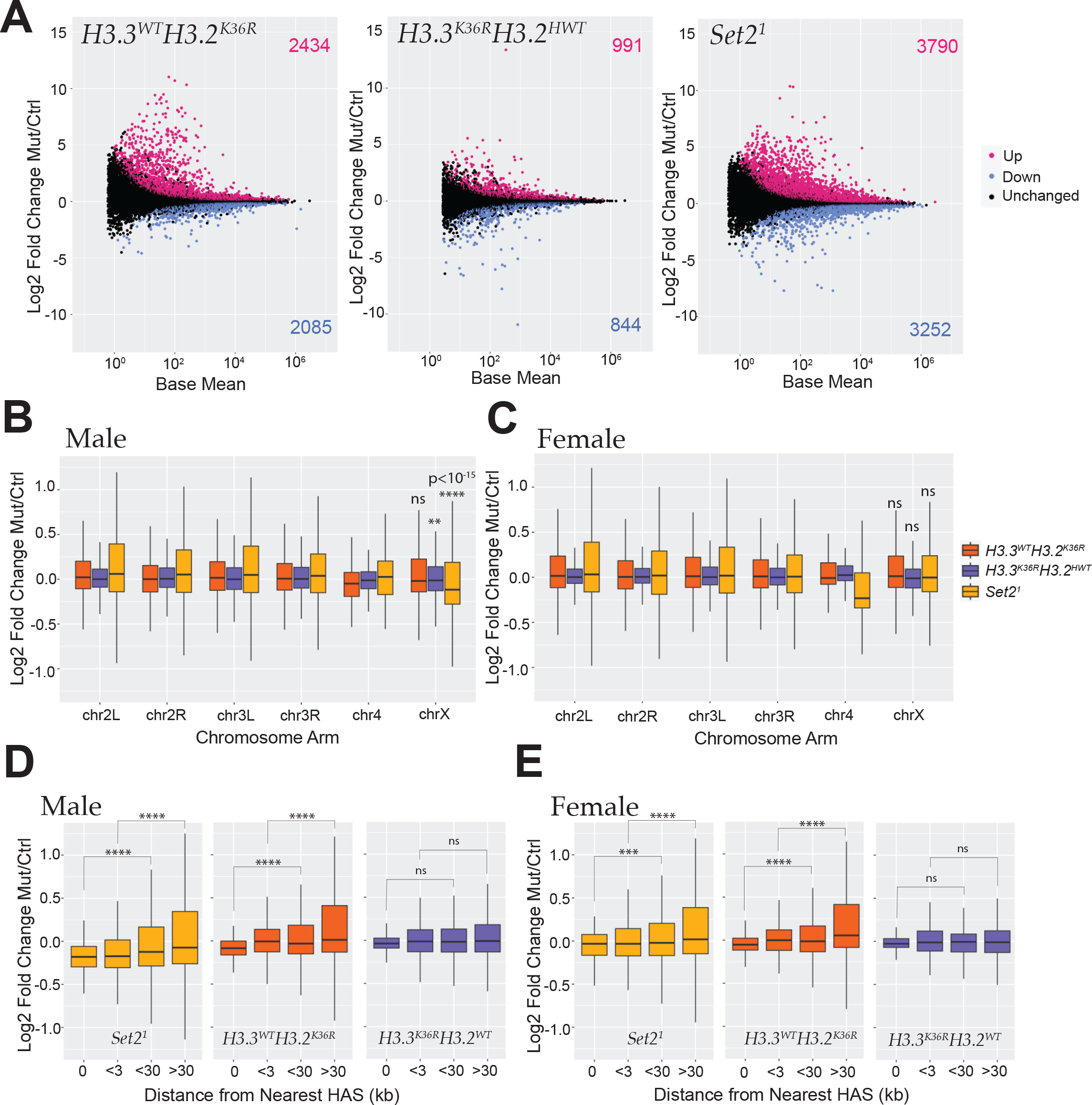
Transcriptomic analyses of dosage compensation in third instar *Set2* and *H3K36R* larval brains. (A) M/A plots comparing gene expression in WL3 brain from combined male and female replicates of mutants relative to control. Mutants represented from left to right with control genotype in parentheses: *H3.3^WT^H3.2^K36R^* (*H3.3^WT^H3.2^HWT^*)*, H3.3^K36R^H3.2^HWT^* (*H3.3^Ctrl^H3.2^HWT^*), and *Set2^1^* (*yw*). Magenta and blue dots represent differentially expressed genes (DEGs) that were significantly (adjusted p-value, p-adj < 0.05) up- or down-regulated, respectively. The number of DEGs in each direction is shown in the upper and lower corners. **(B)** For DESeq2 analyses separated by sex, all genes with a defined P value (not NA), Log2 Fold-change values of mutant genotypes in A, relative to controls were plotted for male replicates and binned by chromosome arm. Median Log2 Fold-change values of X-chromosome genes were compared to the combined set of large autosome (2L, 2R, 3L, and 3R) genes, and p-values computed using the Kruskall-Wallis ANOVA, followed by Dunn’s multiple comparisons tests. \*\**P* < 0.01, \*\*\*\**P* < 10*^-15^*. ns, not significant. **(C)** Same as C, but for female replicates. **(D)** HAS site analysis of mutant males relative to controls. Log2 Fold-change values of *Set2^1^* mutant males were plotted, binned by distance from chrX HAS sites defined previously [31]. \*\*\**P* < 0.001. \*\*\*\**P* < 0.0001. ns, not significant. **(E)** Same as D, but for female replicates.

### Individual Set2 or H3K36 mutations exert weak and inconsistent effects on global X chromosome gene expression

To understand the extent to which H3.3K36, H3.2K36, and Set2 might play a role in DC, we performed additional DESeq2 comparisons, this time separated by sex (Fig.S7). Overall patterns of gene expression were similar to the combined analysis when separated in this manner (Fig.S7). To gain insight into whether, expression of X chromosome genes is inhibited in *H3.3^K36R^H3.2^HWT^* mutants, we plotted the LFC of each mutant genotype relative to its control, binned by chromosome arm for both males and females (Fig. 2B,C). In line with previous work [39], we observed a significant decrease in chrX gene expression in male *Set2^1^* mutants. Importantly, we did not see this effect in females indicating that this X chromosome-wide decrease is male-specific (Fig. 2C). We also observed a very slight, but statistically significant decrease (adjusted *P* < 0.01) in the *H3.3^K36R^H3.2^HWT^* mutant males (Fig. 2B). No change was observed in *H3.3^K36R^H3.2^HWT^* females, or in either sex in the *H3.3^WT^H3.2^K36R^* genotype (Fig. 2B,C). These results suggest that there must be either functional compensation between H3.3K36 and H3.2K36 with respect to male X chromosome gene expression, or that Set2 regulates male X gene expression via some other target. Remarkably, we also observed strong sexual dimorphism in the effect of all three mutant genotypes with respect to the 4^th^ chromosome, implying that sex differences in chromosome-wide gene expression may not always be due to dosage compensation (Fig. 2B,C).

One feature of reduced expression of MSL complex members is a change in the severity of male X gene expression impairment that varies by distance from high-affinity MSL binding sites (HASs; see [31, 57]). Impairment of MSL2 binding to HAS loci results in the greatest degree of gene expression loss overlapping the site itself, whereas impairment of MSL3 exhibits the opposite pattern with the greatest decrease farthest from HAS sites [31, 57]. These and other findings suggest that MSL2 is required for initiation of MSL mediated DC and that MSL3 is involved in spreading of the complex to surrounding genes (reviewed in [14, 58]). We were curious if the small, but significant decrease in X-gene expression in *H3.3^K36R^H3.2^HWT^* males would exhibit an HAS distance trend, consistent with a role in DC. We also wanted to examine whether the previously observed relationship between HAS site distance and gene expression in *Set2^1^* mutants [39] was male-specific.

To probe these questions, we performed HAS distance analysis in both male and female *Set2^1^, H3.3^WT^H3.2^K36R^*, and *H3.3^K36R^H3.2^HWT^* mutants. As shown previously, we observed the greatest decrease in chrX gene expression nearest to HASs in *Set2^1^* males, suggestive of an initiation defect rather than a spreading defect (Fig. 2D [31, 39, 57]). We also detected a similar, but smaller, effect in female brains (Fig. 2E). Analysis of *H3.3^K36R^H3.2^HWT^* mutants demonstrates gene expression trend related to HAS distance, suggesting that the small difference in male X expression may not be due to DC. Inversely, we observed an overall trend in the *H3.3^WT^H3.2^K36R^* males and females resembling that of *Set2^1^* mutants, though weaker and less consistently. On the whole, these observations call into question whether Set2 is likely to be involved in MSL complex spreading, as the observed effects are neither male-specific, nor do they resemble a situation of impaired MSL3 function. Furthermore, we found no evidence that either H3.2 or H3.3 K36R mutation impacts DC at this developmental stage.

### H3.3K36 exhibits differential effects on X-gene expression during development

Genetic redundancy between H3.2K36 and H3.3K36 complicates a determination of the requirement for H3K36me3 in MSL complex spreading. However, one would expect that compensation between H3 variants might be partially bypassed in tissues or developmental stages where one variant predominates. In the adult brain, cells are largely senescent and H3.3 incorporation increases with age [59, 60]. We therefore, took advantage of *H3.3^K36R^* mutant transcriptomic data obtained in adult male and female heads of both “young” (newly eclosed) and “old” (∼23 days post-eclosion) flies [56]. Indeed, transcriptomic dysregulation on the whole increases in *H3.3^K36R^* mutants with age in brain/head tissue (Fig. 2A, [56]). Of note, *H3.3^K36R^* mutant and *H3.3^Ctrl^* animals were on a genetic background with a wild-type RD histone locus in these analyses from adult heads [56].

Chromosome arm plots of LFC values by age and sex show a larger decrease in median LFC for chrX genes relative to the large autosomes for both young and old flies of both sexes (Fig. 3A,B). The magnitude of decrease increases with age, concurrent with increased H3.3 incorporation (Fig. 3A,B). The presence of this decrease in both sexes suggests this effect is due to “X-ness” rather than to DC. If this were true, one prediction would be that despite decreased global X expression, there would be no relationship between LFC and HAS distance. In fact, we observe no relationship in young males and old females, and a significant *upregulation* of chromosome X genes by HAS distance in young females (Fig. 3C, D). In old males, the overall trend is significant, but does not exhibit a consistent change at each increment as would be expected if *H3.3K36* mutation were impeding MSL complex spreading (Fig. 3D)

**Fig. 3.**
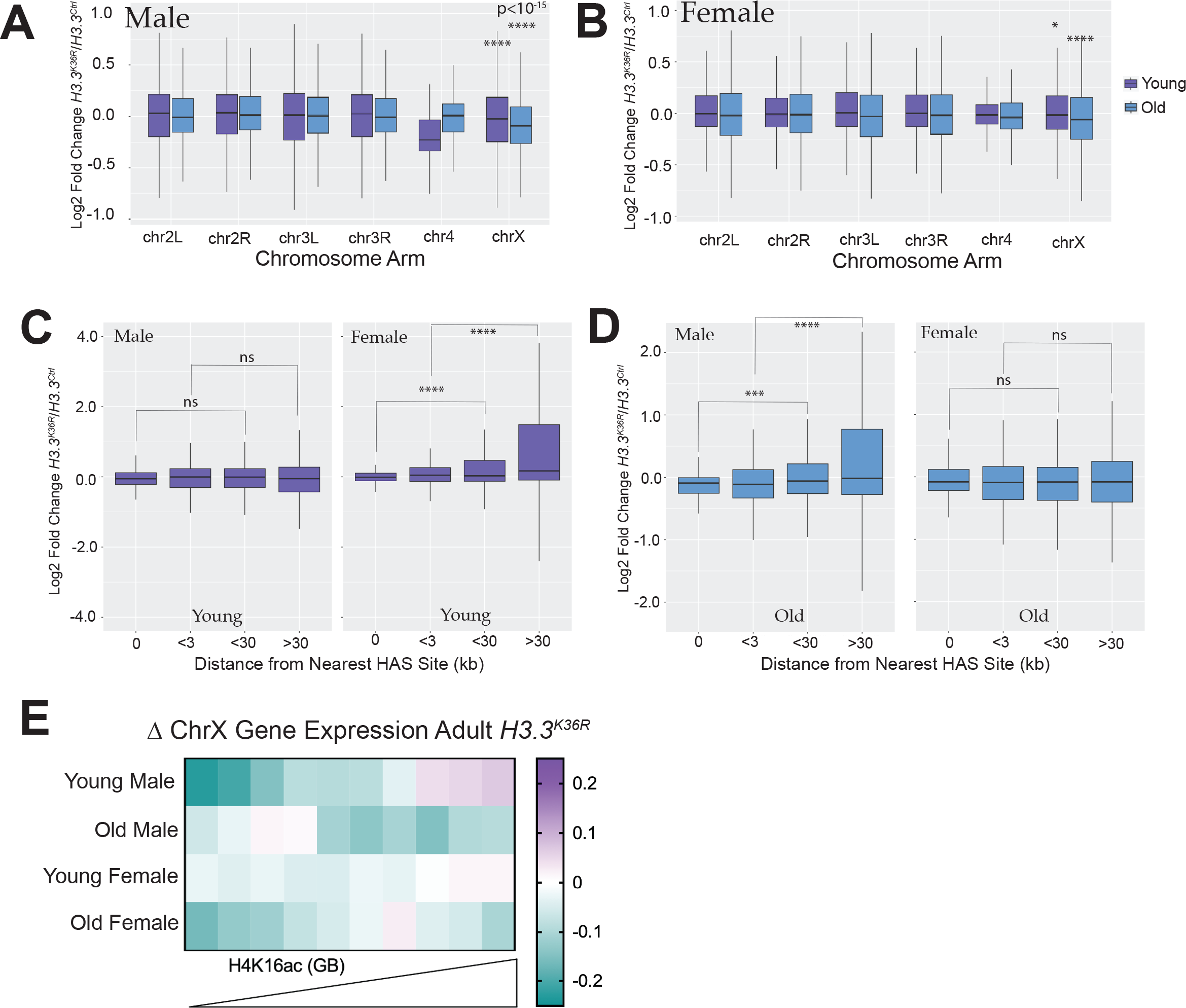
Transcriptomic analyses of dosage compensation in *H3.3^K36R^* adults. (A) For DESeq2 analyses in adult heads separated by sex, and all genes with a defined P value (not NA), Log2 Fold-change values of *H3.3^K36R^* mutants, relative to *H3.3A^null^* controls for young (∼1 day post-eclosion) and old (∼23 day post-eclosion) were plotted for male replicates and binned by chromosome arm. Median Log2 Fold-change values of X-chromosome genes were compared to the combined set of large autosome (2L, 2R, 3L, and 3R) genes, and p-values computed using the Kruskall-Wallis ANOVA, followed by Dunn’s multiple comparisons tests. \**P* < 0.05, \*\*\*\**P* < 10*^-15^*. **(B)** Same as A, but for females. **(C)** HAS site analysis of mutant males relative to controls. Log2 Fold-change values of *H3.3^K36R^* mutant males were plotted, binned by distance from chrX HAS sites defined previously [31]. \*\*\**P* < 0.001. \*\*\*\**P* < 0.0001. ns, not significant. **(D)** Same as D, but for female replicates. **(E)** For ChrX genes, median Log2 Fold-change values for each group were binned by mean H4K16ac ChIP-seq signal in gene bodies from male adult heads and plotted on a heatmap .

**Fig. 4.**
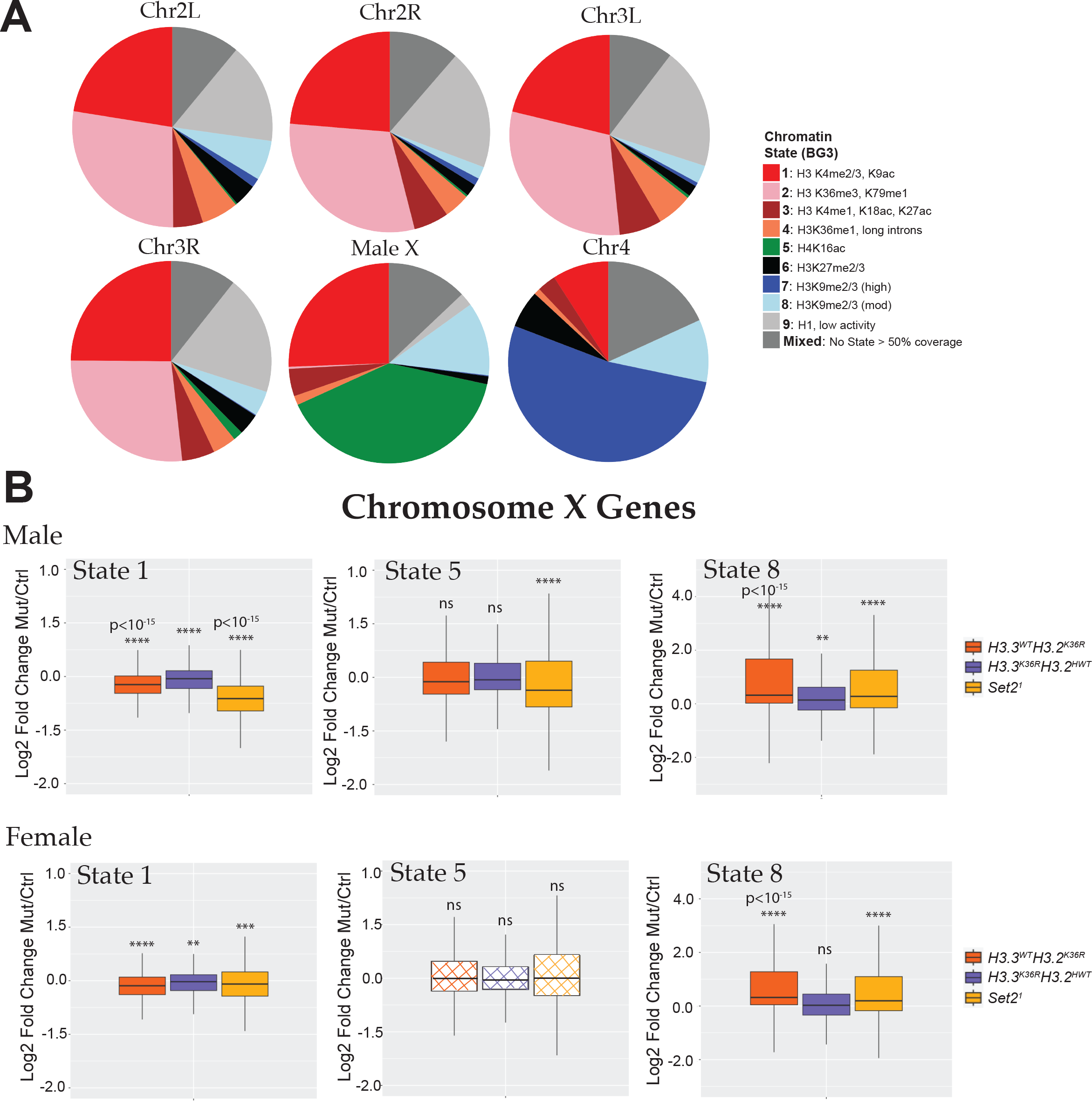
Chromosome X genes with Different Predominant Chromatin States Respond Differently to *Set2* and *H3K36R* mutation. (A) Pie charts depicting predominant chromatin states (defined in [62]) of six *Drosophila* chromosomes in BG3 cells. BEDtools was used to assign genes to a predominant chromatin state. Genes were binned to a given state if > 50% of the gene was marked by that state. Genes where no state color was > 50% of gene length were designated as “Mixed”. Representative histone marks in each state depicted in the legend. A full characterization of each state is described in the source publication [62]. **(B)** Log2 Fold-change values of mutant genotypes described in Fig. S5 for ChrX genes were plotted separately for genes in the three predominant states on the male X: states 1 (n=499), 5 (n=798), and 8 (n=239). Statistical significance of difference between medians was assessed using the Wilcoxon signed rank test, followed by theBenjamini- Hochberg False Discovery Rate (FDR) correction for multiple comparisons. \*\**P* < 0.001. \*\*\**P* < 0.001. \*\*\*\**P* < 0.0001. ns, not significant.

Finally, if H3.3K36me3 promotes DC in aged male flies, we would expect to observe the greatest decreases in X-gene expression on genes with the highest levels of H4K16ac. To assess the relationship between gene expression change and H4K16ac, we binned chrX genes by mean H4K16ac signal in adult heads and plotted LFC in these bins (Fig. 3E).

Unexpectedly, in young male *H3.3^K36R^* fly heads, we observed a compelling, male-specific trend whereby gene expression *increases* with increasing H4K16ac (Fig. 3E). This is precisely the opposite of what one would expect if H3.3K36me3 enables MSL3 spreading. Instead, this pattern is more consistent with H3.3K36 *inhibiting* DC in some way. Also unexpectedly, this relationship changes in the ageing male flies where the genes in the top six deciles of H4K16ac exhibit decreased expression (Fig. 3E). This effect is mirrored (but to a lesser extent) in females (Fig. 3E). These data argue against a simple role for H3.3K36me3 in mediating MSL complex spreading, and instead hint that the effect of H3.3K36 on X-gene expression may be mediated by other processes. Furthermore, these data imply that effects of H3.3K36 on chrX gene expression are influenced by developmental stage and age.

### The effect of Set2 and H3K36 mutations on X genes depends on chromatin context

The effects of *Set2^1^, H3.3^WT^H3.2^K36R^*, and *H3.3^K36R^H3.2^HWT^* mutations on global X chromosome expression neither track consistently by sex, nor do they exhibit predicted trends in gene expression by proximity to HASs. These findings suggest that such effects are unlikely to be caused by a defect in MSL spreading. Furthermore, the largest effect in *Set2^1^* mutant males is considerably weaker than that observed following depletion of MSL complex proteins, and stands in marked contrast to effects in *H4^K16R^* mutants [19, 31, 57].

Given that all chromosomes harbor genes within different chromatin environments, subject to different modes of regulation and activity [61, 62], we wondered whether our observations could be explained by heterogeneous responses to Set2/K36 mutation within different chromatin compartments.

To investigate this hypothesis, we utilized the genome-wide chromatin characterization model defined by Kharchenko and colleagues [62]. This study applied a machine learning approach to ChIP-seq data to define 9 basic chromatin states in two cell culture models. We used their BG3 model (derived from male WL3 larval brain) for this analysis. The 9 chromatin states include 5 “active” states (1-5) and 4 “repressive” states (6- 9). Though most genes span multiple states, we were able to identify a “predominant” chromatin state for most genes, defined as the state covering > 50% of gene body length (Fig. 3A). When genes were classified in this way, the composition of the male X was clearly different from the autosomes, with three states comprising the bulk of genes (Fig. 3A).

State 5 genes, marked by H4K16 acetylated chromatin, encompass nearly half of the genes on the male X. State 1, marked by H3K4me3 and H3K9ac and common at active promoters accounts for about ∼25%. Lastly, repressive State 8, marked by moderate levels of H3K9 di- and trimethylation, covers ∼12% of genes.

To examine whether Set2, H3.2K36 and H3.3K36 regulat chromosome X genes heterogeneously within different chromatin states, we next plotted WL3 brain LFC values of chrX genes for each mutant and sex binned by predominant state (Fig. 3B). Of note, because BG3 cells are male, the chromatin features of these “State 5” genes in females are unknown, but unlikely to be characterized by genic H4K16ac since this is a hallmark of male DC. Remarkably, we observe different patterns of effects in the three mutant genotypes depending on chromatin state (Fig. 3B). For State 5 genes, we observe a significant median decrease specifically in *Set2^1^* males, and no change in *Set2^1^* females or the *H3^K36R^* mutants. However, we note that a substantial fraction (>25%) of State 5 genes are actually upregulated in *Set2^1^* mutant males. In contrast, State 1 genes exhibit significantly reduced expression in both sexes for all three mutant genotypes. This difference reveals that State 5 and State1 chrX genes are differentially sensitive to H3K36 mutation. Even so, the median decrease in expression of State 1 genes in *Set2^1^* males is substantially greater than for the other genotypes (∼6 fold > than *Set2^1^* females; ∼2.5 >*H3.3^WT^H3.2^K36R^* males). The disproportionate effect in both active states in *Set2^1^* males demonstrates that Set2 enhances expression of active genes on the male X in a distinctive manner. Whether this outsized effect is due to an alternative function of Set2 or redundancy between H3.2K36 and H3.3K36 at these genes remains unclear.

In contrast, expression of genes in repressive State 8 are substantially *increased* in *Set2^1^* and *H3.3^WT^H3.2^K36R^* mutants of both sexes, and slightly in *H3.3^K36R^H3.2^HWT^* males. This adds to mounting evidence implicating H3K36 in repressing inactive of lowly expressed genes [56, 63], and implies that that Set2 may support gene repression in some contexts as well. Taken together, these data hint that the effects of Set2, H3.2K36, and H3.3K36 on chrX gene expression are context-dependent.

### Set2 and H3K36 variants exhibit variable patterns of X chromosome gene regulation

Thus far, our analyses hint that chrX genes respond in a pleotropic manner to mutation of *Set2, H3.2K36,* and *H3.3K36*, suggesting that regulation by these players is context-dependent, and potentially multi-faceted. We wanted to better understand the interplay of these mutations on specific genes and genomic contexts, and ascertain whether any of these contexts were associated with sexually dimorphic effects. To address these questions, we first identified groups of genes likely to be similarly regulated. We reasoned that genes with common regulatory mechanisms would exhibit similar patterns of expression changes with respect to genotype and sex. To assess global patterns of regulation across differentially expressed genes on the X, we constructed a *k*-means clustered heatmap of the combined DEGs for all mutants. We used the *z*-score difference of DESeq2 normalized counts (individual replicates – mean of controls of combined sexes) for each gene to enable comparison of genes with vastly different expression levels (Fig. 5A). From this heatmap, we were able to extract gene names for further analysis of cluster features. For each cluster, we calculated the base mean gene expression (Fig. 5B), LFC between mutants and same-sex controls (Fig. 5C), relative levels of H3K36 methylation states (Fig. 5D) and DC proteins (Fig. 5E), and relative enrichment of proteins and marks associated with the Kharchenko chromatin states (Fig. S8). For analyses of cluster features, chrX genes unchanged in any of the Set2/H3K36 mutants (nonDEGs) were included for comparison. Of interest, this *k-*means clustering approach reveals that many X-genes exhibit mild sexual dimorphism in expression in wild type males and females (Fig. 5A), as male and female replicates are consistently on opposite sides of the genotype mean (L3-c1, c2, c3, c4, c9) in the *yw* control (Fig. 5A).

**Fig. 5.**
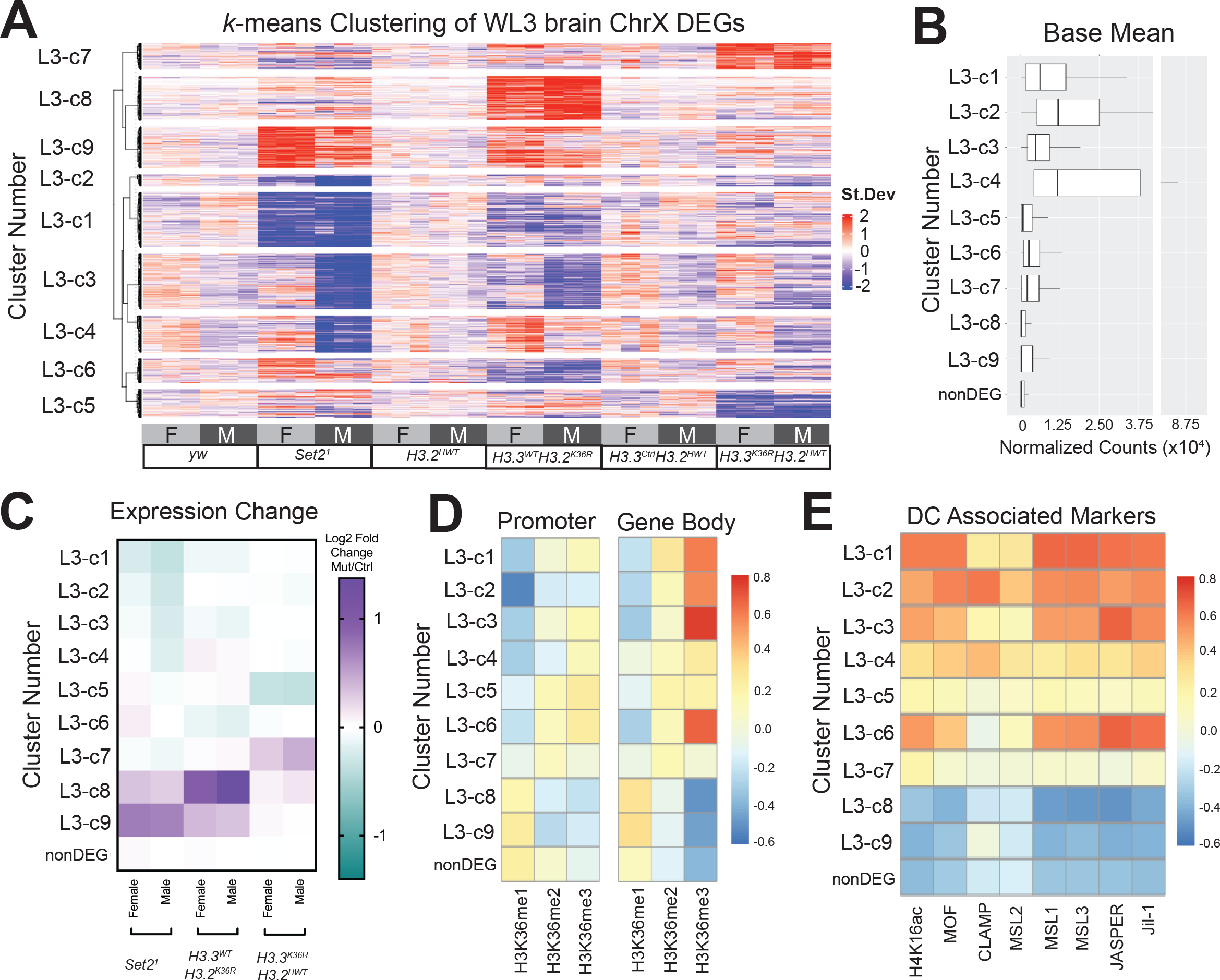
***k*-means clustering of gene expression reveals heterogeneous regulation of Chromosome X genes by *Set2*/*H3K36R*. (A)** DESeq2 normalized count values for the combined set of Chromosome X DEGs from WL3 brain from all mutant and control genotypes were *z*-score normalized by gene to put all expression values on the same scale (mean=0, SD=1). Differences of individual replicate values for each gene were calculated relative to the mean *z*-score of control replicates. A *k*-means clustered heatmap (*k*=9) was generated from these values. Below the heatmap, genotype and sex of each replicate is indicated. To the left, cluster numbers for subsequent analyses are indicated. N values for each cluster are as follows: c1 (272), c2 (57), c3 (275), c4 (180), c5 (141), c6 (120), c7 (132), c8 (219), c9 (205), not differentially expressed (nonDEG, 416). **(B)** Base Mean of normalized DESeq2 counts for DEGs (binned by Cluster Number in panel A) and Non-DEGs (genes not differentially expressed in any mutant genotype). **(C)** Chromosome X genes were grouped by gene expression cluster (Panel A), and a heatmaps of median Log2 Fold-change values for mutant genotypes, separated by sex, was constructed. **(D)** Mean levels of H3K36me1, me2, and me3 in BG3 cells were calculated for genes from all chromosomes for both promoter and gene body regions. For each methyl state, *z*-scores were computed for all genes. For X-genes, median *z*-score was computed for Panel A heatmap clusters and non-DEGs, and a heatmap of these values was constructed to highlight relative levels of H3K36 modification states between clusters. **(E)** Relative abundance of DC modifications and proteins within gene bodies were calculated and plotted as in (D), using datasets generated in S2 cells.

With respect to our genotypes of interest, we identified nine distinct patterns of regulation amongst all genotypes and sexes, three of which (clusters L3-c1, c2, c3; 604/2017 of total chrX genes) align with what would be expected if H3K36me3 enabled spreading of the MSL complex (Fig. 5A,C). For these clusters, we observed male-specific expression decreases in the *Set2^1^* mutant, and to a lesser extent in either the *H3.3^WT^H3.2^K36R^* or the *H3.3^K36R^H3.2^HWT^* mutants (Fig. 5C). These clusters were also amongst the highest in relative enrichment of H3K36me3 and MSL complex proteins (Fig. 5D, E). Notably, we did not observe any gene clusters with expression changes in the *Set2^1^* mutants, but where *H3.3^WT^H3.2^K36R^* and *H3.3^K36R^H3.2^HWT^* resembled controls, suggesting that the role of *Set2^1^* in promoting expression of chrX genes in males is likely to occur by way of H3K36 in this tissue/stage, rather than by some other target or function of Set2. L3-c1,c2, c3 are compatible with the idea of redundancy between variants, as the magnitude of change in the *Set2^1^* mutant is greater than either *H3^K36R^* mutant even while changing in the same direction (Fig. 5C). These observations are consistent with the possibility that Set2 via H3K36me3 may promote gene expression of some dosage-compensated genes.

Two other clusters also exhibited sexually dimorphic expression changes, but different from what would be expected if H3K36me3 were facilitating canonical DC. Cluster L3-c4 shows decreased expression in *Set2^1^* males, but *increased* expression in *H3.3^WT^H3.2^K36R^* females, whereas L3-c6 shows *increased* expression in *Set2^1^* females and decreased expression in *H3.3^WT^H3.2^K36R^* males (Fig. 5A,C). L3-c6 is among the most enriched in H3K36me3 in gene bodies and L3-c4 is relatively less so. Increased expression in female mutants resembles what would be predicted in response to a defect in “non-canonical dosage compensation” whereby lowly expressed genes in heterochromatin depleted of MSL complex in males, are inhibited in females by way of homolog pairing [8]. However, neither cluster is depleted in MSL complex proteins (Fig. 5E) or enriched in repressive histone marks or chromatin proteins (Fig. S8). Furthermore, L3-c4 contains genes with the highest base mean (Fig. 5B). These observations suggest that L3-c4 and L3- c6 are unlikely to employ non-canonical DC as defined previously.

Clusters L3-c7, c8, c9 are primarily defined by upregulation in one or more mutant genotype. L3-c8 and L3-c9 are relatively enriched in H3K36me1 and depleted in H3K36me3 (Fig.5D). These genes were lowly expressed on the whole and enriched in heterochromatic marks (Fig. 5B, S8). Even so, gene expression was significantly increased in L3-c9 in the *Set2^1^* mutant (Fig.5C). This is consistent with the possibility of indirect effects, or these genes may correspond to genes where Set2 depletion results in increased H3K36me1 on the chromosome arms [47]. Lastly, clusters L3-c5 and L3-c7 are driven primarily by H3.3 mutation. These genes also have intermediate levels of DC proteins. Overall, these data imply a large degree of heterogeneity in how H3.2K36, H3.3K36, and Set2 impact X chromosome gene expression, which is inconsistent with a role in chromosome-wide dosage compensation.

### Insulator proteins associate with X chromosome DEGs in Set2/H3K36 mutants

We next wanted to gain insight into what might be driving the diverse patterns of gene expression changes observed in the *Set2* and *H3^K36R^* mutants. To this end, we performed motif enrichment analysis using the SEA (Simple Enrichment Analysis) tool [64] on the WL3 brain mRNA-seq heatmap clusters (Fig 5A; Fig 6A) [65]. Promoter and gene body regions for genes in each cluster were compared to these regions in nonDEGs. We focused on the most enriched motifs, those exhibiting a *q*-value < 0.05 and enrichment over control sequences > 2 (Fig. 6A).

**Fig. 6.**
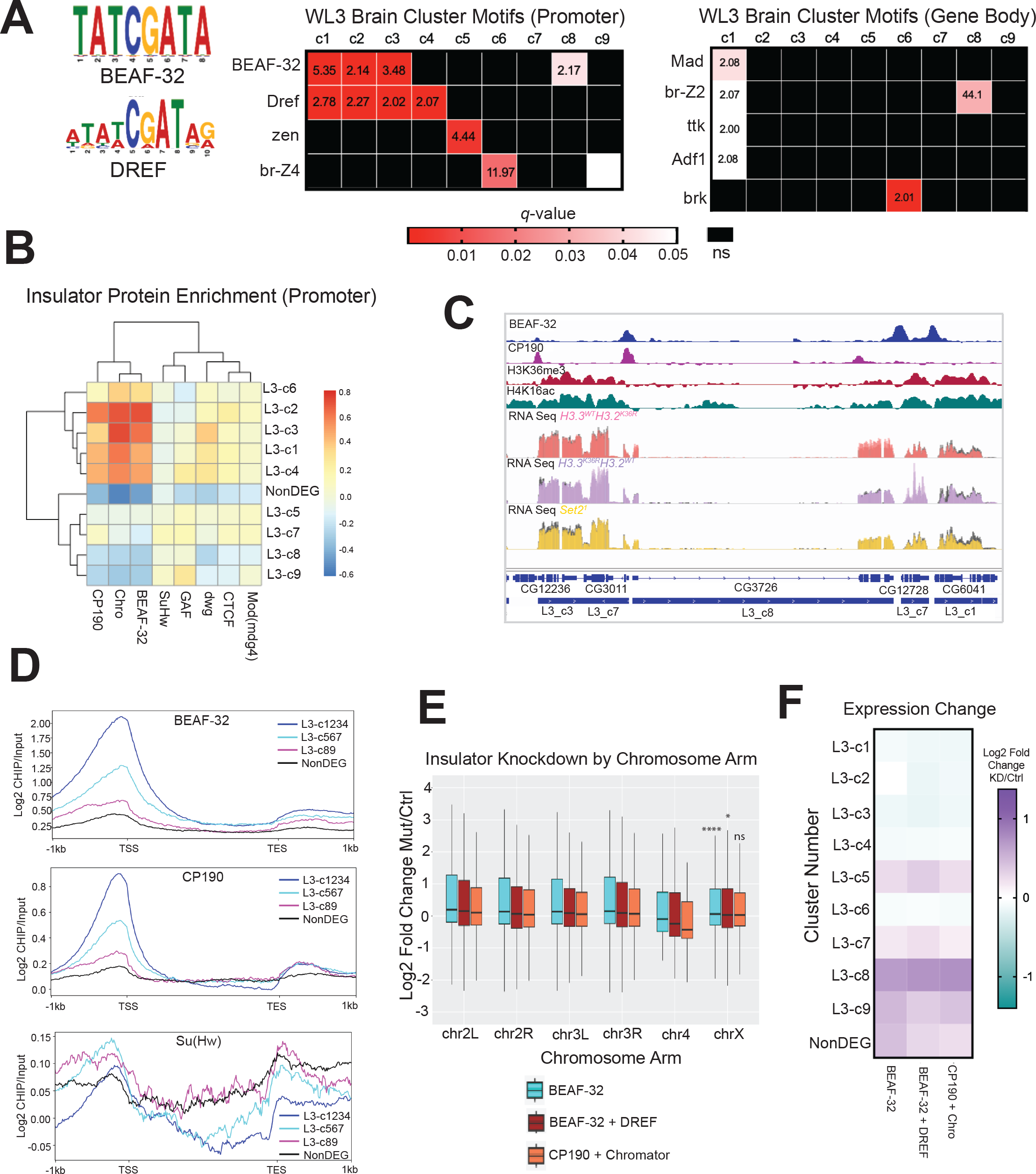
Motif analysis reveals that Set2/K36 and BEAF-32 regulate common gene sets similarly. (A) SEA (Simple Enrichment Analysis) was performed on promoters and gene bodies of gene groups from Fig. 5A using motifs from the FLYREG.v2 database (Fig 5A; Fig 6A [64, 65]). Motifs with *q-*value > 0.05 and enrichment value over nonDEG control sequences > 2 were displayed. Motifs not meeting either threshold were designated as ns and colored black. **(B)** A heatmap of insulator protein binding was generated as in Fig. 5D and 5E, with the addition of hierarchical clustering. **(C)** Browser shot of insulator protein binding relative to H3K36me3, H4K16ac, L3 cluster annotation, and WL3 brain male RNA-seq data. RPGC normalized RNA-seq data is colored for each mutant with control data shown in gray on the same track line. **(D)** Scaled gene metaplots of insulator proteins with genes grouped by similar enrichment of BEAF-32 and CP190. SuHw is included for comparison. **(E)** Chromosome arm plots and accompanying statistical analyses were produced as in Fig. 2B from RNA-seq data from insulator transcript knockdowns in BG3 cells generated by [66]. **F)** Median LFC values for BEAF-32, BEAF-32 + DREF, and CP190 + Chromator were determined for each L3 gene cluster from Fig. 5A and plotted as a heatmap.

Interestingly, BEAF-32 and Dref motifs were enriched at promoters across multiple clusters, and exhibiting diverse expression patterns between mutants (Fig. 6A). BEAF-32 is a protein linked to 3D genome organization, insulator function, and gene regulation [66–69]. Dref is a transcription factor involved in insulator function, chromatin organization, gene expression, and telomere maintenance [66, 70, 71]. Interestingly, BEAF-32 and Dref bind similar, often overlapping, DNA motifs [72]. Both functional redundancy [66, 73] and inverse binding profiles have been reported for these factors in different contexts [72]. The most significantly enriched clusters for Dref motifs (L3-c1, L3-c2, L3-c3, and L1-c4), also have the highest median gene expression and exhibit a male-specific decrease in gene expression in the *Set2^1^* mutants (Fig. 5A,B). Three of these clusters (c1, c2, and c3) are also the most significantly enriched in BEAF-32 motifs (Fig. 6A). L3-c8 was also enriched in BEAF-32 motifs, though these genes were upregulated in *Set2*/*H3^K36R^* mutants (Fig. 5A,B).

Next, we assessed whether motif enrichment corresponded to increased insulator protein binding at the promoters of these genes. We constructed heatmaps of relative insulator protein binding for each L3 heatmap cluster for factors with available modENCODE ChIP data (as in Fig.5D,E). We included proteins known to work in conjunction with BEAF-32 (CP190 and Chromator) along with others that operate in different insulator complexes (SuHw and GAF) [69]. We observed substantial relative enrichment of BEAF-32, CP190, and Chromator in L3-c1, c2, c3, and c4 (Fig. 6B). Of note, L3-c8 was relatively depleted in binding of these proteins, despite enrichment of BEAF-32 motifs (Fig. 6B). We observed peaks of BEAF-32 and CP190 at many promoters and some 3’ ends of genes, but these peaks did not always overlap with each other (Fig.6C). For comparison, we saw no enrichment of SuHw on any cluster or the NonDEGs (Fig. 6B).

We also constructed metaplots of BEAF-32 and CP190 to assess the distribution of signal across genes with similar levels of binding (Fig. 6D). Consistent with previous reports, BEAF-32 and CP190 peak near the TSS, with a much smaller enrichment after the TES ([74]; Fig. 6D). This effect was strongest in L3-clusters 1-4, and weakest in the nonDEGs (Fig. 6D). In contrast, a metaplot of SuHw showed relative depletion in L3- clusters 1-4 (Fig. 6D).

The male X chromosome of BEAF-32 mutants exhibit unusual morphology in polytene spreads, despite normal recruitment of MOF [75]. Tissues and cells with impaired levels of BEAF-32 also have widespread transcriptomic changes [66, 76]. We wondered whether cells with a reduction in BEAF-32 might exhibit a decrease in chrX gene expression relative to autosomes, as was observed in *Set2^1^* mutant males ([39]; Fig. 2B). To address this question, we reanalyzed RNA-seq data from a previous study of BG3 cells RNAi depleted for insulator complex transcripts [66]. We calculated LFC values for knockdown (KD) conditions of BEAF-32, BEAF-32 + Dref, and CP190 + Chromator and plotted these values by chromosome arm (Fig. 6E).

Like *Set2^1^* mutant males, median gene expression for autosomal genes was elevated for all three insulator KD conditions (Fig. 2B, 6E). Expression of chrX genes was also elevated in the insulator KD conditions, but for the BEAF-32 and BEAF-32 + Dref conditions, this increase in expression was significantly less than what was observed in autosomes (Fig. 6E). In contrast, there was no significant difference in the CP190 + Chromator condition between autosomes and chrX, despite ∼90% and ∼70% reductions in CP190 and Chromator proteins, respectively (Fig. 6E, [66]. These data imply that BEAF-32 promotes gene repression to a lesser degree on the male X chromosome than on autosomes.

Given the heterogeneous, context-dependent effects on chrX gene expression when components of the Set2/H3K36 axis are mutated, we wanted to determine if reduction of insulator components demonstrated similarly heterogeneous changes. We hypothesized that if Set2/K36 and BEAF-32 dependent mechanisms of gene regulation were operating on the same genes in a collaborative manner, one would observe similar gene expression trends in BEAF-32 knockdown cells when binned according to Set2/H3K36 expression clusters. When this analysis was performed, we observed a remarkable concordance between the gene expression trends in the *Set2^1^* mutant males and the insulator protein knockdowns for nearly all L3 clusters (Fig. 5C, 6F). The exceptions were L3-c5 and L3-c7 which were primarily driven by changes in the *H3.3^K36R^H3.2^HWT^* mutant. In summary, these data demonstrate that BEAF-32 binds the promoters of Set2 responsive chrX genes in male cells, and that mutation of both factors have similar effects on expression of dosage-compensated genes. This is consistent with the possibility that Set2 and H3K36 may enhance expression of many male X genes by impacting insulator function rather than by way of MSL complex spreading.

### H3K36me3 does not play an essential role in MSL3 spreading

Our experiments thus far suggest that H3K36me3 is unlikely to be uniquely important for MSL complex spreading. For chrX genes, mutation of *Set2* in males causes small decreases in downregulated genes, and upregulates many others. Moreover, many of the same changes can be observed in females to a lesser extent. In some gene groups, the effects of *Set2* and *H3^K36^* mutation do not align. These effects are consistent with the possibility that the Set2/H3K36 axis is affecting gene expression by one or more other means, including by impacting insulator function. However, recent work suggests that MSL3 might also bind H3K36me2, which could explain the weak and inconsistent effect on chrX gene expression in the *Set2^1^* mutants [47]. Furthermore, we have not yet fully investigated the prospect of functional redundancy between H3.2K36 and H3.3K36.

To address these alternatives, we performed total RNA-seq and DESeq2 analysis at the L1 stage in *Set2^1^* and combined *H3.3^K36R^H3.2^H3K36R^* mutants where all zygotic H3K36 has been mutated, alongside control genotypes. The *H3.3^K36R^H3.2^K36R^* genotype addresses both genetic redundancy between variants and the possibility that MSL3 might bind to H3K36me2, simultaneously. We used a mixed sex population because sexing them at this stage in the context of a transgenic system already using YFP selection was not yet possible. Because we used mixed sex larvae, we also included the *H3.3^Ctrl^H4^K16R^* mutant genotype to verify that we could detect a signature of male DC in a mixed sex population. We examined this developmental stage because the *H3.3^K36R^H3.2^K36R^* mutants are L1 lethal [51].

Genome-wide MA plots of *Set2^1^* and *H3.3^K36R^H3.2^K36R^* mutants illustrate that large numbers of genes are differentially expressed in both mutants (6,533 and 5,799 respectively), comparable to that observed in *Set2^1^* mutant WL3 brains, indicating that maternal contribution of wild type proteins is unlikely to be masking an effect on gene regulation (Fig. 7A, 2A). In contrast, a modest number (645) of genes reached statistical significance in the *H3.3^Ctrl^H4^K16R^* mutants. These overall trends were preserved when a cutoff of LFC > |1| was employed for these DEGs (Fig. S9A). Despite the relatively small number of DEGs in the *H3.3^Ctrl^H4^K16R^* animals, when we plotted LFC values by chromosome arm, there was a highly significant (*p* < 10^-15^) decrease in global chrX gene expression in these animals, demonstrating the ability to detect a DC defect in a mixed population (Fig. 7B). In contrast, despite much greater changes to their respective transcriptomes, we observed no change in the *Set2^1^* mutants and a highly significant *increase* in the *H3.3^K36R^H3.2^K36R^* genotype (Fig. 7B).

**Fig. 7.**
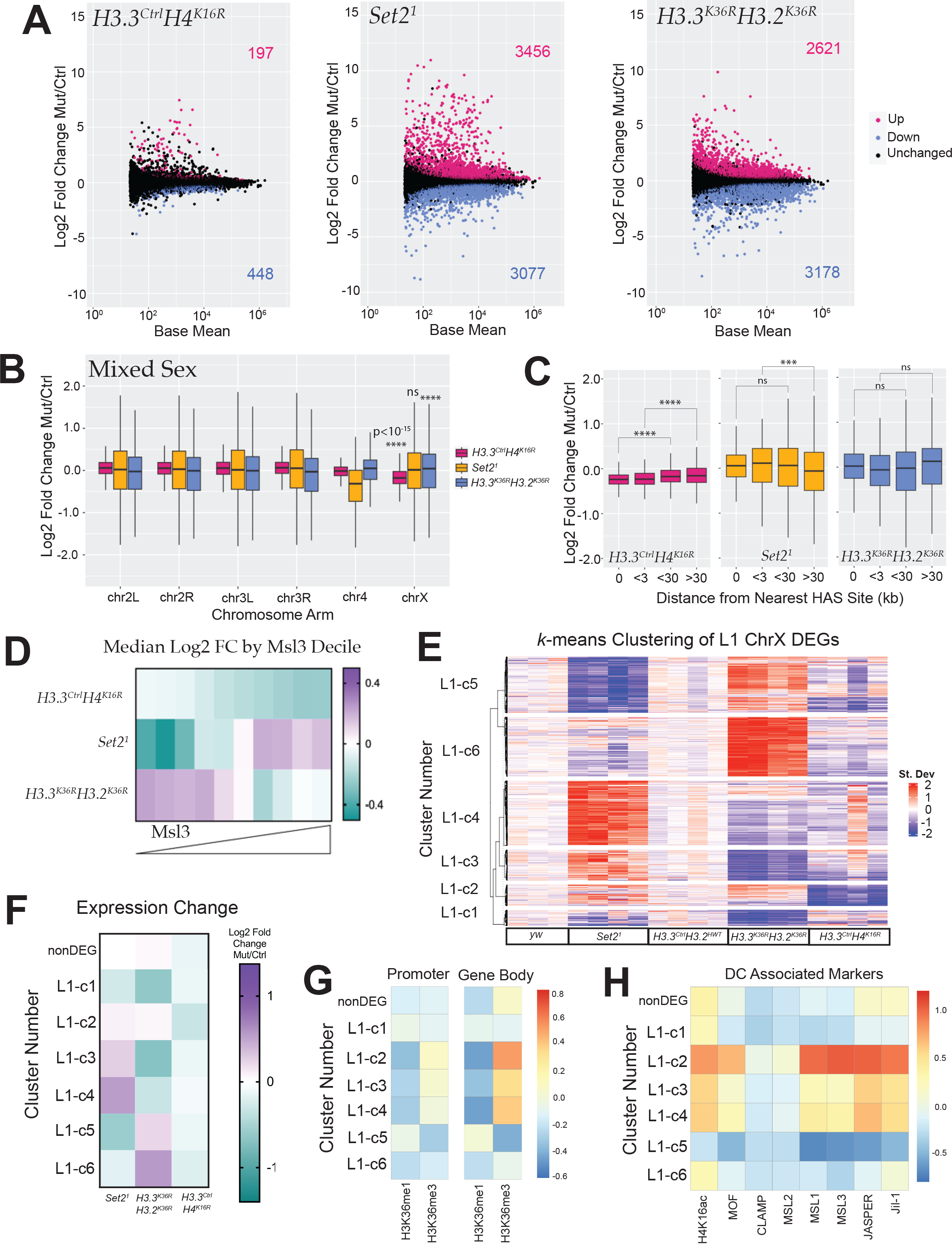
Transcriptomic analyses of dosage compensation in first instar *Set2* and combined *H3K36R* larvae. (A) M/A plots comparing gene expression changes from mixed sex, whole L1 animals. Mutants represented from left to right with control genotype in parentheses: *H3.3^Ctrl^H4^K16R^* (*H3.3^Ctrl^H3.2^HWT^*)*, Set2^1^* (*yw*), and *H3.3^K36R^H3.2^K36R^* (*H3.3^Ctrl^H3.2^HWT^*). Magenta and blue dots represent differentially expressed genes (DEGs) that were significantly (adjusted p-value, p-adj < 0.05) up- or down-regulated, respectively. The number of DEGs in each direction is shown in the upper and lower corners. **(B)** For all genes with a defined P value (not NA), Log2 Fold-change values of mutant genotypes in A, relative to controls were plotted and binned by chromosome arm as in Fig.2B,C. **(C)** HAS site analysis of mixed sex animals relative to controls was analysed as in Fig. 2D,E. **(D)** Mean MSL3 levels across gene bodies in S2 cells (see Methods). ChrX genes were binned by MSL3 decile, and heatmaps of median Log2 Fold-change values for mutant genotypes were constructed. **(E)** A *k*-means clustered heatmap (*k*=6) was generated as in Fig.5A (see Methods). Genotypes are indicated below heatmap; clusters to the left. N values for each cluster are as follows: c1 (163), c2 (125), c3 (182), c4 (385), c5 (333), c6 (353), not differentially expressed (nonDEG, 456). **(F)** ChrX genes were grouped by gene expression cluster (Panel E), and a heatmap of median Log2 Fold-change values for mutant genotypes, was constructed. **(G)** Heatmaps of median H3K36me1, me2, and me3 per cluster were constucted as in Fig. 5E, except using ChIP data from S2 cell. **(H)** Relative abundance of DC modifications and proteins within gene bodies were calculated and plotted as in (G).

HAS distance analyses were concordant with these results. In the *H3.3^Ctrl^H4^K16R^* mutants, we observed clear and statistically significant incremental change in the magnitude of transcript reduction varying by distance from the HAS site (Fig. 7C).

Conversely, we found no such correlation in the *Set2^1^* and *H3.3^K36R^H3.2^K36R^* mutants (Fig. 7C).

If H3K36 methylation were required for MSL complex spreading, one prediction would be that the greatest loss of expression would be on genes with the most MSL complex. To test this, we plotted median LFC for decile bins corresponding to mean gene body MSL3 signal (Fig. 7D). In the *H3.3^Ctrl^H4^K16R^* mutant controls, we observed a nearly perfect incremental relationship between bin medians whereby the greatest decrease in gene expression occurs at the highest MSL3 levels (Fig. 7D). This trend was clearly visible in mixed sex samples and with relatively less transcriptome dysregulation overall. In contrast, the *Set2^1^* mutants tended to *increase* at genes with the highest MSL3 (Fig. 7D). In the *H3.3^K36R^H3.2^K36R^* genotype, there was little or no change in the top two deciles of MSL3 occupancy, with the most substantial median decrease in gene expression occurring in the fourth highest decile (Fig. 7D). Importantly, neither the *Set2^1^* or *H3.3^K36R^H3.2^K36R^* genotype showed any clear relationship with MSL3 occupancy. Nor did those two genotypes resemble one another in this aspect. Instead, they trended opposite to each other in all but one bin (Fig. 7D). These opposite trends also held when LFC values were binned by base mean gene expression (Fig. S9B).

Next, we wanted to look directly at the patterns of gene expression among *Set2^1^*, *H3.3^K36R^H3.2^K36R^,* and *H3.3^Ctrl^H4^K16R^* mutants for genes on the X chromosome. We constructed a *k*-means clustered heatmap of *z*-score differences for the combined set of chrX DEGs, as in Fig. 5A (Fig. 7E). Strikingly, we observed that most genes exhibit an opposite expression trend between the *Set2^1^* and *H3.3^K36R^H3.2^K36R^* mutants, providing further evidence of a regulatory “switch” between methylation states (Fig. 7E, F). We also observed that the cluster with the strongest decrease in expression in the *H3.3^Ctrl^H4^K16R^* mutants (L1- c2), the highest relative H3K36me3 (Fig. 7G), and greatest relative occupancy of DC related proteins (Fig. 7H), showed a trend toward *upregulation* in both the *Set2^1^* and *H3.3^K36R^H3.2^K36R^* mutants, which argues against a role for H3K36me3 in promoting H4K16ac (Fig. 7E). Furthermore, the three clusters with the highest relative enrichment of H3K36me3 (L1-c2, c3, c4), show *upregulation* in the *Set2^1^* mutant suggesting that Set2 is acting to dampen expression at these genes (Fig. 7F,G).

### Set2 and H3K36 exhibit context-specific expression discordance

We also noted that *k-*means clustered heatmaps looked very different at L1 and L3 stages (Fig. 5A, Fig.7E). In the L3 heatmap, *Set2^1^* and *H3^K36R^* mutations resulted in only 3 of the 9 clusters (L3-c4, c6, and c7, comprising ∼27% of L1 DEGs) exhibiting discordant expression changes (Fig.5A,C). In contrast, for nearly all gene clusters in the L1 heatmap, *Set2^1^* and the combined *H3.3^K36R^H3.2^K36R^* mutant resulted in opposite trends (∼81% of L1 DEGs), excepting L1-c1 and L1-c2 (Fig. 7E,F). In the case of the *Set2^1^* mutant, we also see discordance between developmental stage/tissue type within the very same genotype.

Analyses of the 3 most common male X Kharchenko states (Fig. S9C; States 1, 5, and 9 in S2 cells) reveals contradictory trends in State 1 for this genotype (Fig. 4B, S9D). This reveals an additional layer of context-dependence in X chromosome regulation related to developmental stage or tissue type. Intriguingly, the relative levels of the three *Drosophila* H3K36 methyltransferases can also differ between WL3 brain and whole L1 larvae, consistent with the possibility that differential methylation profiles at particular loci could mediate these changes (See Discussion) (Fig. S10). In summary, these data provide compelling evidence for context- and stage-dependent regulation of the X chromosome by Set2/H3K36. Moreover, the data do not support a requirement for a specific H3K36 methylation state in MSL complex spreading, even when all zygotic copies of H3 cannot be methylated at lysine-36.

## Discussion

### Trimethylation of H3K36 is not essential for spreading of the MSL complex

This study provides strong evidence against the prevailing dogma that H3K36me3 mediates spreading of the MSL complex. Although many gene clusters enriched in MSL complex members in males are downregulated in *Set2* mutant males, most of these genes exhibit the same general trends in females (Fig. 5A,C, E). Furthermore, we have identified genes marked by MSL and highly decorated with H3K36me3 that are unaffected in *Set2* males, but trend upwards in *Set2* females (Fig. 5, L3-c6). *H3.3A^null^H4^K16R^*We also note that HAS analyses of *Set2* mutants resemble the pattern observed in depletion of MSL2 (involved in initiation at HASs) rather than MSL3 (involved in MSL complex spreading ([31, 57]; Fig. 2D,E). *H3.3^Ctrl^H4^K16R^* mutants, even at an early stage and in a mixed sex population, exhibit a nearly ubiquitous downward trend in chrX gene expression (Fig.7). In contrast, mutations of Set2 and H3K36 elicit heterogeneous effects across the X chromosome at multiple developmental stages (Figs 4, 5, and 7).

Yet, clearly for a large proportion of genes exhibiting enrichment of H4K16ac and MSL complex, Set2 exerts an outsized effect in males (Fig. 5A,C,E). We propose a model whereby Set2 (via H3K36) likely supports expression of genes by other mechanisms such as nucleosome turnover [77], elongation control [78–80], recruitment of HDACs [81, 82], or as suggested in this study, functional relationships with insulator proteins (Fig. 6; [67, 83])).

In males, one or more of these mechanisms may synergize with the MSL complex, which is believed to utilize both elongation control and 3D genome organization in propagating its function [12, 49, 57, 84].

If H3K36me3 is not essential for MSL complex spreading, what are some alternatives? One possibility is methylation of histone H4 lysine 20 (H4K20). Like H3K36me3 and H4K16ac, H4K20 monomethylation localizes preferentially to gene bodies [85–87]. *In vitro* studies demonstrate that H4K20me1 and H4K20me2 peptides have an up-to 50fold higher affinity for the MSL3 chromodomain compared to H3K36me3 [88–90]. A Y31A mutation in the MSL3 chromodomain that weakens *in vitro* binding of H4K20 methylated peptides, also reduces survival of males when introduced *in vivo* [90]. The K9- S10 portion of the H3 tail has also been connected to regulation of male X genes. H3K9me2 on X-specific 1.688^X^ satellite sequences has been shown to support proper expression of surrounding genes [91], and ectopic expression of siRNA from these repeats can partially rescue *roX1roX2* mutant males [92].

Importantly, these possibilities are not mutually exclusive. MSL complex might make use of multiple chromatin features for targeting, including H3K36me3, H4K20me, and H3K9me2. This could occur either redundantly between marks, or with specificity on a gene-by-gene basis depending on which marks predominate. The second possibility might be evidenced by preferential regulation of different subsets of male X genes in H3K36, H4K20, H9K9 mutants. There is precedent for redundancies in the DC system regarding both *roX1* and *roX2*, as well as replication-dependent *H4^K16^* and replication-independent *His4r* [19, 20]. Further studies addressing the impact of these other histone tail residues on DC, either alone or in concert, would be informative.

### Relationships between H3K36, insulator proteins, and dosage compensation

Given that we found enrichment of BEAF-32 and CP190 in the promoters of Set2 responsive X-genes (Fig. 6A,B), and similar effects on many gene clusters when Set2 and BEAF-32 are impaired (Figs. 5C & 6F), we believe that 3D genome structure and insulator function are especially promising areas of potential synergy between H3K36 and DC. The male and female X chromosomes have surprisingly similar large-scale organization [49, 93], but with more mid- to long-range interactions on the male X [94]. Intriguingly, Clamp, a protein essential for *Drosophila* DC [32, 33, 95, 96] promotes the interaction of HASs in 3D space [97]. Furthermore, Clamp and MSL complex binding are enriched at BEAF-32/CP190 domain boundaries that are weakened in males [94]. Like H3K36me3, Clamp binds genome-wide where it can impact gene expression independently of the MSL complex, as well as synergize with the MSL complex during DC [96, 98–100]. Thus, Clamp sets a precedent for the model that we espouse.

Interestingly, Clamp is known to interact with with two separate insulator complexes: the late boundary complex [101] and the *gypsy* insulator [102]. Furthermore, depletion of Clamp results in reduction of CP190 at some sites [102]. Clamp has also been show to interact with two separate insulator complexes: the late boundary complex[101] [102] [102]Clamp also [33, 99]interacts with several histone proteins, including H3.2 and H3.3 [103], and can bind nucleosomal DNA to increase chromatin accessibility [98]. Thus, it is tempting to speculate that H3K36 and Clamp may cooperate in some manner.

BEAF-32 peaks occur most often near the TSS, while H3K36me3 is enriched at the 3’ ends of genes, thus any model of interplay between these factors must account for their different spatial positions. One possibility is an interaction between BEAF-32 and H3K36me3 chromatin. Indeed, one 4C study identifying the most prevalent chromatin states for BEAF-32 interactions showed that BEAF-32 had the strongest interaction with active chromatin harboring H3K36me3, rather than active chromatin depleted of H3K36me3, consistent with the possibility of a functionally important interaction [61, 68]. One study reports that weakening of domain boundaries containing BEAF-32 parallels binding of the MSL complex on the male X [94]. In conjunction with our data, this suggests the intriguing possibility that H3K36me3 might assist in weakening these boundaries somehow. Future 4C or Hi-C studies, as well as chromatin binding studies of BEAF-32 and other insulator proteins in *Set2* and *H3^K36R^* mutants would be of great interest in evaluating this hypothesis.

### Context-dependence of X-gene expression at different developmental stages

One surprising conclusion of our study is the strong effect of developmental stage/tissue type on X chromosome gene expression heterogeneity. We enumerate two distinct effects. First, we find that the degree of agreement between Set2 and H3K36 mutants differs widely between the L1 and WL3 brain datasets, with much greater discordance in the L1 samples (Fig. 5A,C & Fig. 7E, F). Secondly, we find that individual genotypes can trend differently in the same chromatin states between these datasets. The best example of this is in the *Set2^1^* mutant genotype in State 1 (Fig.4B, Fig.S9D).

What could be causing these variations? One exciting possibility is that differential expression of H3K36 methyltransferases (KMTs) at different stages or in different tissues could be driving these differences. In our RNA-seq data, we see distinct relative levels of H3K36 KMTs between L1, WL3 brain, and adult head (Fig. S10). At L1, NSD and Ash1 are ∼40% and ∼15% more highly expressed than Set2 (Fig. S10). In contrast, NSD is ∼15% more highly expressed than Set2 in WL3 brain, while Ash1 expression falls below that of Set2.

In adult heads, NSD expression is less than 50% of that of Set2 and Ash1, which are roughly equal (Fig. S10). Some of these differences may be specific to nervous system tissue, as another study examined levels of these KMTs and found different trends in whole WL3 larvae and whole aged adults [39].

One model driven primarily by experiments in female Kc cells posits a direct interaction between BEAF-32 and NSD which preconditions H3K36me2 for Dref/Set2 driven trimethylation [67, 83]. Bulk modifications by H4K16ac by Western blot elicited the conclusion that decrease of H3K36me3 alone leads to decreased H4K16ac, while decrease of both H3K36 di- and trimethylation led to *increased* H4K16ac [83]. Since *H3^K36R^* mutation eliminates all methylation states while *Set2* mutation eliminates only trimethylation, this is consistent with the idea of a regulatory switch between methylation states, and could account for some of the discordance we observed, while also explaining how these differences could be exacerbated by varying levels of H3K36 KMTs. It is also intriguing to speculate that given this connection with insulators, differential KMT levels might also exert differential effects on insulator function.

Though interesting, this “preconditioning model” has recently been challenged by a genome-level study in S2 cells of the three *Drosophila* H3K36 methyltransferases (KMTs), their binding patterns, and the subsequent effects on H3K36 methylation and the transcriptome when these writer enzymes are subjected to RNAi knockdown [47]. This study suggests that Set2 does not require H3K36me2 to trimethylated H3K36, and that most genes are primarily methylated by one particular KMT on a gene-by-gene basis [47]. Even so, reduction of one KMT can also affect activity of other KMTs in a “see-saw effect” [47]. The authors also report that NSD can perform trimethylation on some genes. One possible implication of this study is that differential levels of KMTs would be expected to exert genome-wide, locus-specific, and context-dependent effects that could conceivably vary by tissue and/or developmental stage. A comprehensive investigation of H3K36 readers and writers in different cell types, tissues, and stages would shed additional light on the basis for these context-dependent effects.

Although we believe we can make many strong conclusions, it is important to point out potential limitations of this study. First, these results are limited to specific developmental timepoints/tissues. While we would expect findings related directly to MSL complex function to be broadly applicable, other sources of heterogeneity are likely to vary in other tissues and stages, as we have found to be the case in this study. The use of mixed sex larvae at L1, while suggestive, necessitates cautious interpretation. ChIP-Seq datas were obtained from cell culture models. Additionally, we have not directly measured MSL3 binding, but have inferred it by examining gene expression. In future studies, we would like to generate antibodies to test this directly.

## Conclusions

In summary, the work here does not support the widely held view [21, 34, 104–106] that H3K36me3 is essential for *Drosophila* MSL complex spreading. Our transcriptomic study of X-gene regulation in *Set2*, *H3.2^K36^*, *H3.3^K36^* and combined *H3^K36^* mutants of both sexes is inconsistent with this idea. Instead, the data point to mechanisms whereby Set2 and H3K36 support X chromosome gene expression via processes common to both sexes, that synergize with the MSL complex in males. These findings lead to a more accurate understanding of the relationship between H3K36 writers and residues and its effects on the activity of MSL complex. As these same regulatory paradigms and processes are conserved in mammals, these findings will be important for our understanding of human health and disease.

## Methods

### *Drosophila* lines and husbandry

To obtain experimental progeny, parental flies were housed in cages sealed with grape juice agar plates smeared with supplemental yeast paste. Plates were changed daily. L1 larvae were obtained directly from the grape juice plates. Older animals were picked at the L2 stage, 50 per vial, and raised on cornmeal-molasses food. All experimental animals were raised at 25°C. Details concerning construction of BAC transgenes generated previously containing the 12xH3.2 and 12xH4K16R histone gene arrays can be found in [19, 53, 107].

*His*Δ indicates *Df(HisC^ED1429^)*; flies containing the *HisΔ*, *twGal4*, and *HisΔ*, *UAS:2xYFP* chromosomes [108] were received from A. Herzig. The *H3.3A^2x1^ (H3.3A^null^)* [109], *Set2^1^* allele and rescue transgene [84], *Df(2 L)Bsc110* deficiency, and the beta-tubulin GFP protein trap stock used for recombination with the rescue transgene were obtained from Bloomington Stock Center (nos. 68240, 77917, 8835, and 50867). The *H3.3B^K36R^* CRISPR allele was generated previously [51]. Gene names, annotations, genome sequence, references, and other valuable information useful to this study were acquired from FlyBase [110].

### Generation of mutant genotypes

For detailed genetic schemes, see Figs. S1, S2, S4, & S5). *HisΔ* animals were obtained by selection for yellow fluorescent protein (YFP). Other H3.3 genotypes were selected for absence of a *CyO*, *twGFP* balancer chromosome. Set2 mutants were detected by absence of GFP from both a maternal FM7i, act>GFP balancer and a paternal chromosome carrying a Set2 rescue transgene linked to a transgene expressing GFP tagged Β-tubulin.

### Pupal and adult viability and sex ratio assays

For each genotype, fifty L2 larvae were picked from grape juice agar plates and transferred to vials containing molasses-cornmeal food. Full plates were picked to prevent bias due to different developmental timing between males and females. Pupae and eclosed adults were counted until 13 and 18 days after egg laying, respectively. Pupal and adult eclosion percentages were calculated per-vial by dividing the number of pupal cases or eclosed adults per 50 input larvae and multiplying by 100. Each vial constituted one biological replicate for statistical purposes. Between 400 and 500 total animals (8-10 replicate vials) were analyzed per genotype. For male and female ratios, number of males and females were determined from eclosed adults from the above viability assays. Statistical significance for % eclosion was obtained with Brown-Forsythe and Welch ANOVA tests, followed by Dunnett’s T3 multiple comparisons test. Statistical significance for sex ratio was obtained with Fisher’s Exact Test, followed by the Benjamini-Hochberg False Discovery Rate (FDR) correction for multiple comparisons (Q=0.05). Graphpad Prism was used for calculations.

### RNA Seq library preparation and sequencing

For the wandering L3 brain experiment, 25 brains were dissected per replicate and homogenized in 1ml Trizol solution. RNA was obtained from the Trizol aqueous phase using the Zymo RNA Clean and Concentrator-5 kit (Genesee Scientific #11-352) plus DNAse I treatment, according to manufacturer’s instructions. PolyA-selected libraries were prepared using the KAPA stranded mRNA kit (Roche # 07962207001) and sequenced using the NOVASeq-S1 paired-end 100 platform. For the L1 experiment, 25-30 larvae were picked, rinsed with PBS, homogenized in 1mL Trizol, and isolated above. Total RNA Seq libraries were prepared with Nugen Ovation Universal Drosophila kit and sequenced with NOVASeq-S4 paired-end 100 platform.

### Bioinformatic analyses

For both sequencing experiments, reads were trimmed for adaptor sequence/low-quality sequence using BBDuk (bbmap). FastQC was used for quality control [111], and reads were aligned to genome build DM6 using the STAR aligner [112]. Aligned reads were counted with featureCounts [113] and differential expression analyses were completed with DESeq2 [114]. Of note, for the L1 data, one genotype (*H3.3^K36R^H3.2^HWT^*) from the same sequencing run was included in construction of the DESeq model, but not included in any downstream analysis. *k*-means clustered heatmaps of *z*-score differences from RNA Seq data were produced as follows. The combined set of chromosome X DEGs for all mutant genotypes were used for each heatmap. *z*-scores for each gene were obtained from DESeq2 normalized counts for each replicate. For each gene, *z*-score differences were obtained by: *z^replicate^* – *z^mean_ctrl_reps_both_sexes^*. For each *z*-score difference, the mean of the most appropriate control genotype was used. Scree plots were used to determine the value of *k.* The ComplexHeatmap package was used to plot *z*-score differences [115]. Gene lists for each cluster were exported for downstream analyses of cluster features. Boxplots were made using *ggplot2* from the Tidyverse package [116]. Heatmaps displaying median LFC values per bins of MSL or H4K16ac were made using GraphPad Prism for Mac, GraphPad Software, www.graphpad.com. Heatmaps displaying median *z-*scores of ChIP Seq data per RNA Seq cluster were produced as follows. For modENCODE data files, DM3 aligned bedGraph files were converted to bigwig files using Crossmap [117]. For H3K36me2 ChIP, data was downloaded from SRA, and sequences were trimmed, quality checked, and aligned as above. BAM files from ChIP files were normalized to input files and output to bigwig format using deepTools [118]. For Clamp, MSL2, MSL3, Jasper, and Jil-1 RNAi data generated by previously, DM6 aligned bigwigs were downloaded directly from the GEO repository [28, 32]. BEDTools was used to calculate mean ChIP signal over promoter regions (500bp upstream of the TSS) and gene bodies for each gene [119]. *z*-scores for mean promoter and gene body ChIP signal were obtained relative to all chrX genes. For each heatmap of median ChIP Seq signal values (Figs. 5D, 5E, 6B, 7G, 7H, SXX) for RNA Seq gene clusters generated in Fig. 5A & Fig. 7E, a median *z*-score for each cluster for each ChIP dataset was calculated and plotted using the pheatmap package [120]. *z-*score normalization enabled relative comparisons between different histone modification or chromatin binding protein datasets obtained using different antibodies and conditions.

Motif analysis was performed by the SEA (Simple Enrichment Analysis) tool using a predefined set of motifs [65]. Metaplots were generated from modENCODE ChIP data for genes in each RNA heatmap cluster using deepTools [118]. Browser tracks for genomic data were visualized on the Integrated Genomics Viewer (IGV) [121].

Statistical analyses for RNA-seq data is as follows. Significant DEGs were determined by DESeq2 with and adjusted *p*-value <0.05. For chromosome arm plots, LFC values of X-chromosome genes were compared to the combined set of large autosome (2L, 2R, 3L, and 3R) genes, and *p*-values computed using the Kruskall-Wallis ANOVA, followed by Dunn’s multiple comparisons tests. For predominant chromatin state analyses based on [62], Statistical significance of the difference between medians was obtained using the Wilcoxon signed rank test and theBenjamini-Hochberg False Discovery Rate (FDR) multiple comparisons correction.

## Acknowledgements

We thank the UNC High Throughput Sequencing Facility (HTSF) for their assistance with generating the datasets used here, and members of the Matera laboratory for helpful discussions and critical reading of the manuscript. This work was supported by the National Institutes of Health (NIGMS) grant R35-GM136435 (to A.G.M.).

