## Supplementary Information for "Set2 and H3K36 regulate the *Drosophila* male X chromosome in a context-specific manner, independent from MSL complex spreading"

#### Supplementary Figures and Legends

**<u>Fig. S1. Crossing schemes and genotype info for Panels 1B&1C.</u>** Crossing schemes used to generate the sex genotypes assayed in Figure 1B and 1C. Maternal and paternal genotypes are indicated above the boxes containing experimental progeny. Both full and abbreviated genotypes are listed for experimental animals.

**<u>Fig. S2. Crossing schemes and genotype info for Figures 3 & S3.</u>** Crossing schemes used to generate the sex genotypes assayed in Figure 1B and 1C. Maternal and paternal genotypes are indicated above the boxes containing experimental progeny. Both full and abbreviated genotypes are listed for experimental animals.

**<u>Figure S3. Sex ratio of -$H3.3^{K36R}$ and $H3.3^{Ctrl}$ animals.</u>** Proportion of male and female eclosed animals were calculated from viability experiments from animals in Figs 3&S2. Statistical significance for sex ratio was calculated with GraphPad Prism software using Fisher's Exact Test. ns, not significant.

**<u>Fig. S4. Crossing schemes and genotype info for Panels 1D&1E.</u>** Crossing schemes used to generate the sex genotypes assayed in Figure 1B and 1C. Maternal and paternal genotypes are indicated above the boxes containing experimental progeny. Both full and abbreviated genotypes are listed for experimental animals.

**<u>Fig. S5. Crossing schemes and genotype info for Figures 2, 4, 5, 6, & 7.</u>** Crossing schemes used to generate the sex genotypes assayed in Figures 2,4,5,6&7. Maternal and paternal genotypes are indicated above the boxes containing experimental progeny. Both full and abbreviated genotypes are listed for experimental animals. $Set2^1$ mutants were selected by GFP- status,unlike the histone mutants,which were YFP+.Note, $Set2^1$ mutants were selected by GFP- status,unlike the histone mutants,which were YFP+.

**<u>Fig. S6 Additional analyses of WL3 brain DESeq2.</u> A)** Principle Component Analysis (PCA) of WL3 brain RNA-seq data set. **B)** To the left, a Pie chart depicting the number of genes with defined *p*-values and the overlap of significant DEGs between genotypes. To the right, for the subset of genes differentially expressed in all three mutant genotypes, a pie chart depicting the most common patterns of change within this category. **C)** MA plots employing a LFC cutoff of > |1| for the combined-sex DESeq2 analysis. For genes meeting this cutoff, DEGs with and adjusted *p*-value > 0.05 are colored magenta for upregulated genes; blue for downregulated genes. The number of genes in each category is indicated in those same colors.

**<u>Fig. S7 Sex-specific MA plots.</u>** M/A plots comparing gene expression in WL3 brain from DESeq2 analyses separated by sex. Mutants represented from left to right with control genotype in parentheses: $H3.3^{WT}H3.2^{K36R}$ ($H3.3^{WT}H3.2^{HWT}$), $H3.3^{K36R}H3.2^{HWT}$ ($H3.3^{Ctrl}H3.2^{HWT}$), and $Set2^1$ (*yw*). Magenta and blue dots represent differentially expressed genes (DEGs) that were significantly (adjusted p-value, p-adj < 0.05) up- or down-regulated, respectively. In the case of plots C&D, an additional LFC > |1| was met for coloration. The number of DEGs in each direction is shown in the upper and lower corners. **(A)** Plots of male data, adjusted p-value, p-adj < 0.05. **(B)** Plots of female data, adjusted p-value, p-adj <

0.05. **(C)** Plots of male data, adjusted p-value, p-adj < 0.05 & LFC > |1|. **(D)** Plots of male data, adjusted p-value, p-adj < 0.05 & LFC > |1|.

**Fig. S8 Relative binding of histone modifications/proteins for chromatin states 1-9.** Mean levels of histone modifications and chromatin binding proteins characteristic of the 9 chromatin states described previously (See Figs.4,5) were calculated for genes from all chromosomes for both promoter and gene body regions. For each datasetChIP*z*-scores were computed for dataset, *z* Fig.5A and chrX -scores were computed for chrX*z*-score was computed genes. For each cluster from Fig.5A and chrX Below, histone modifications and proteins are color-coded according to the primary chromatin state that it most represents. A state surrounded by a red box indicates that this mark/protein is depleted in that state. Coloration in the secondary line indicates that the mark/protein is also conspicuously high/low in an additional state.nonDEGs, median *z*-score was computed. A heatmap of these values was constructed to highlight relative levels of each modification or chromatin binding protein per cluster. Below, histone modifications and proteins are color-coded according to the primary chromatin state that it most represents. A state surrounded by a red box indicates that this mark/protein is depleted in that state. Coloration in the secondary line indicates that the mark/protein is also conspicuously high/low in an additional state.

**Fig. S9 Additional plots for L1 RNA-seq.** **(A)** M/A plots comparing gene expression changes from mixed sex, whole L1 animals. Mutants represented from left to right with control genotype in parentheses: $H3.3^{Ctrl}H4^{K16R}$ ($H3.3^{Ctrl}H3.2^{HWT}$), $Set2^{1}$ (*yw*), and $H3.3^{K36R}H3.2^{K36R}$ ($H3.3^{Ctrl}H3.2^{HWT}$). Magenta and blue dots represent differentially expressed genes (DEGs) that were significantly (adjusted p-value, p-adj < 0.05) and LFC >|1|, up- or down-regulated, respectively. The number of DEGs in each direction is shown in the upper and lower corners. **(B)** For chrX genes, Heatmap of LFC of mutants/ctrl for three mutant genotypes binned by increasing Base Mean from DESeq2 output. **(C)** Pie charts depicting predominant chromatin states (defined in [62]) of six *Drosophila* chromosomes in S2 cells. BEDtools was used to assign genes to a predominant chromatin state. Genes were binned to a given state if > 50% of the gene was marked by that state. Genes where no state color was > 50% of gene length were designated as "Mixed". Representative histone marks in each state depicted in the legend. A full characterization of each state is described in the source publication [62]. **(D)** $Log_2$ Fold-change values of three mutant genotypes described in Fig.S5 for ChrX genes were plotted separately for genes in the three predominant states on the male X: states 1 (n=544), 5 (n=762), and 9 (n=232). Statistical significance of difference between medians was assessed using the Wilcoxon signed rank test. followed by theBenjamini-Hochberg False Discovery Rate (FDR) correction for multiple comparisons. **P* < 0.05. ***P* < 0.001. *****P* < 0.0001. ns, not significant.

**Fig. S10 Relative H3K36MT levels from L1, WL3 brain, and adult head.** For Ash1, NSD, and Set2, base mean from DESeq analyses from three datasets used in this study (L1, WL3 brain, and adult head) were normalized to gene length to obtain an estimate of transcript abundance. To account for different read depths for each dataset, Ash1, NSD, and Set2 are presented as a ratio relative to the Set2 level.

**Table S1. Experimental and control genotype pairs.** Experimental genotypes are listed alongside the control genotype used. Control genotypes were selected to have the most similar genetic background to experimental genotypes in terms of Δ*His* and *H3.3A* status. Some control genotypes with the same alleles are designated by multiple names depending on the experimental genotypes in a given experiment. This is denoted by "=". Also listed are figures relevant to each genotype pair.

**Table S2. Publicly available datasets used in this study.** All publicly available datasets are listed in this table with the following information: 1) Relevant histone modification, binding protein, mutation, or knockdown 2) Type of dataset 3) Gene Expression Omnibus (GEO) accession number 4) Cell line, tissue, or stage, and 5) Relevant figure panels. GEO accession numbers are color-coded by source as follows: Black (modENCODE), Red [56], Blue [32], Green [28], and Purple [66].

### Crosses for Panels 1B & 1C

♀ HisΔ, twGal4 / CyO ; $12xH3.2^{HWT}$ / $12xH3.2^{HWT}$ X ♂ HisΔ, UAS:2xYFP / CyO ; $12xH3/H4^{HWT}$ / $12xH3/H4^{HWT}$

**HWT/HWT** = HisΔ, twGal4 / HisΔ,UAS:2xYFP ; $12xH3^{HWT}/H4^{HWT}$ / $12xH3^{HWT}/H4^{HWT}$

♀ HisΔ, twGal4 / CyO ; $12xH3.2^{K36R}$ / TM6B, Tb X ♂ HisΔ, UAS:2xYFP / CyO ; $12xH3^{HWT}/H4^{HWT}$ / $12xH3.2^{HWT}$

**K36R/HWT** = HisΔ, twGal4 / HisΔ,UAS:2xYFP ; $12xH3^{HWT}/H4^{HWT}$ / $12xH3^{HWT}/H4^{HWT}$

♀ HisΔ, UAS:2xYFP / CyO ; $12xH4^{K16R}$ / $12xH4^{K16R}$ X ♂ HisΔ, twGal4 / CyO ; $12xH3^{HWT}/H4^{HWT}$ / $12xH3^{HWT}/H4^{HWT}$

**K16R/HWT** = HisΔ, UAS:2xYFP / HisΔ, twGal4 ; $12xH3.2^{K16R}$ / $12xH3^{HWT}/H4^{HWT}$

♀ HisΔ, UAS:2xYFP / CyO ; $12xH4^{K16R}$ / $12xH4^{K16R}$ X ♂ HisΔ, twGal4 / CyO ; $12xH4^{K16R}$ / $12xH3.2^{K16R}$

**K16R/K16R** = HisΔ, UAS:2xYFP / HisΔ, twGal4 ; $12xH4^{K16R}$ / $12xH4^{K16R}$

♀ HisΔ, twGal4 / CyO ; $12xH3.2^{K36R}$ / TM6B, Tb X ♂ HisΔ, UAS:2xYFP / CyO ; $12xH4^{K16R}$ / $12xH4^{K16R}$

**K36R/K16R** = HisΔ, twGal4 / HisΔ,UAS:2xYFP ; $12xH3.2^{K36R}$ / $12xH4^{K16R}$

♀ HisΔ, UAS:2xYFP / CyO ; $12xH4^{K16R}$ / $12xH4^{K16R}$ X ♂ HisΔ, twGal4 / CyO ; $12xH3.2^{K36R}$ / TM6B, Tb

**K16R/K36R** = HisΔ, UAS:2xYFP / HisΔ, twGal4 ; $12xH4^{K16R}$ / $12xH3.2^{K36R}$

Figure S1

### Crosses for Figures 3 & S3

♀ $\frac{H3.3B^{K36R}\text{ or }+}{H3.3B^{K36R}\text{ or }+}$ ; $\frac{H3.3A^{2x1}}{CyO,\ twiGFP}$ x ♂ $\frac{H3.3B^{K36R}\text{ or }+}{}$ ; $\frac{Df(2L)Bsc110}{CyO,\ twiGFP}$

**$H3.3^{Ctrl}$** = $\frac{+}{}$ ; $\frac{H3.3A^{2x1}}{Df(2L)BSC110}$

**$H3.3^{K36R}$** = $\frac{H3.3B^{K36R.}}{}$ ; $\frac{H3.3A^{2x1}}{Df(2L)BSC110}$

Figure S2

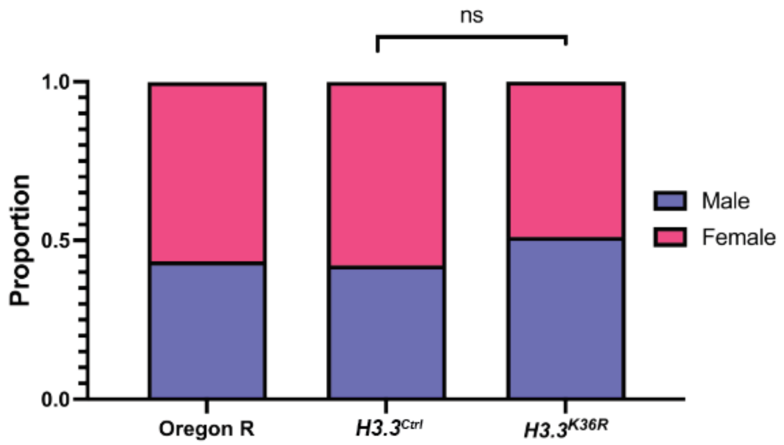


| | | | |
|---|---|---|---|
| *H3.3A* | *WT* | *null* | *null* |
| *H3.3B* | *WT* | *WT* | *K36R* |
| *HisC* | *WT* | *WT* | *WT* |
| *12xTg* | --- | --- | --- |
| (n) | 218 | 321 | 363 |

Figure S3

### Crosses for Panels 1D & 1E

♀ $\frac{+}{+};\frac{His\Delta,\ twiGal4}{CyO}$ X ♂ $\frac{+}{+};\frac{His\Delta,\ UAS{:}2xYFP}{CyO};\frac{12xH3^{HWT}/H4^{HWT}}{12xH3^{HWT}/H4^{HWT}}$

*These two genotypes are equivalent

| | | |
|---|---|---|
| **$H4^{HWT}$** **$H3.3^{WT}H3.2^{HWT}$** | = | $\frac{+}{+};\frac{His\Delta,\ twiGal4}{His\Delta,\ UAS{:}2xYFP};\frac{12xH3^{HWT}/H4^{HWT}}{+}$ |

♀ $\frac{H3.3B^{K36R}\text{ or }+}{H3.3B^{K36R}\text{ or }+};\frac{H3.3A^{2x1},\ His\Delta,\ twiGal4}{CyO}$ X

♂ $\frac{H3.3B^{K36R}\text{ or }+}{};\frac{H3.3A^{2x1},\ His\Delta,\ UAS{:}2xYFP}{CyO};\frac{12xH3^{HWT}/H4^{HWT}\text{ or }12xH4^{K16R}}{12xH3^{HWT}/H4^{HWT}\text{ or }12xH4^{K16R}}$

| | | |
|---|---|---|
| **$H3.3^{Ctrl}H4^{HWT}$** | = | $\frac{+}{+};\frac{H3.3A^{2x1},\ His\Delta,\ twiGal4}{H3.3A^{2x1},\ His\Delta,\ UAS{:}2xYFP};\frac{12xH3^{HWT}/H4^{HWT}}{+}$ |
| **$H3.3^{K36R}H4^{HWT}$** | = | $\frac{H3.3B^{K36R.}}{H3.3B^{K36R}};\frac{H3.3A^{2x1},\ His\Delta,\ twiGal4}{H3.3A^{2x1},\ His\Delta,\ UAS{:}2xYFP};\frac{12xH3^{HWT}/H4^{HWT}}{+}$ |
| **$H3.3^{Ctrl}H4^{K16R}$** | = | $\frac{+}{+};\frac{H3.3A^{2x1},\ His\Delta,\ twiGal4}{H3.3A^{2x1},\ His\Delta,\ UAS{:}2xYFP};\frac{12xH4^{K16R}}{+}$ |
| **$H3.3^{K36R}H4^{K16R}$** | = | $\frac{H3.3B^{K36R.}}{H3.3B^{K36R}};\frac{H3.3A^{2x1},\ His\Delta,\ twiGal4}{H3.3A^{2x1},\ His\Delta,\ UAS{:}2xYFP};\frac{12xH4^{K16R}}{+}$ |

Figure S4

### Crosses for Figures 2, 4, 5, 6, & 7

♀ $\frac{Set2^{1}}{\text{FM7i, act>GFP}} ; \frac{+}{+}$  X  ♂ $\frac{Set2^{1}}{} ; \frac{\{Set2\text{-}Rescue\},\ \{bTub>GFP\}}{CyO}$

**$Set2^{1}$ =** $\frac{Set2^{1}}{} ; \frac{+}{CyO}$

♀ $\frac{+}{+} ; \frac{His\Delta,\ twiGal4}{CyO}$  X  ♂ $\frac{+}{+} ; \frac{His\Delta,\ UAS{:}2xYFP}{CyO} ; \frac{12xH3^{HWT}/H4^{HWT}}{12xH3^{HWT}/H4^{HWT}}$ or $\frac{12xH3.2^{K36R}}{TM6B}$

*These two genotypes are equivalent

**$H3.3^{WT}H3.2^{HWT}$ $H4^{HWT}$** = $\frac{+}{+} ; \frac{His\Delta,\ twiGal4}{His\Delta,\ UAS{:}2xYFP} ; \frac{12xH3^{HWT}/H4^{HWT}}{+}$

**$H3.3^{WT}H3.2^{K36R}$** = $\frac{+}{+} ; \frac{His\Delta,\ twiGal4}{His\Delta,\ UAS{:}2xYFP} ; \frac{12xH3.2^{K36R}}{+}$

♀ $\frac{H3.3B^{K36R}\text{ or }+}{H3.3B^{K36R}\text{ or }+} ; \frac{H3.3A^{2x1},\ His\Delta,\ twiGal4}{CyO}$  X

♂ $\frac{H3.3B^{K36R}\text{ or }+}{} ; \frac{H3.3A^{2x1},\ His\Delta,\ UAS{:}2xYFP}{CyO} ; \frac{12xH3^{HWT}/H4^{HWT}}{12xH3^{HWT}/H4^{HWT}}$ or $\frac{12xH3.2^{K36R}}{TM6B}$

*These two genotypes are equivalent

**$H3.3^{Ctrl}H3.2^{HWT}$ $H3.3^{Ctrl}H4^{HWT}$** = $\frac{+}{+} ; \frac{H3.3A^{2x1},\ His\Delta,\ twiGal4}{H3.3A^{2x1},\ His\Delta,\ UAS{:}2xYFP} ; \frac{12xH3^{HWT}/H4^{HWT}}{+}$

**$H3.3^{K36R}H3.2^{HWT}$** = $\frac{H3.3B^{K36R.}}{H3.3B^{K36R}} ; \frac{H3.3A^{2x1},\ His\Delta,\ twiGal4}{H3.3A^{2x1},\ His\Delta,\ UAS{:}2xYFP} ; \frac{12xH3^{HWT}/H4^{HWT}}{+}$

**$H3.3^{K36R}H3.2^{K36R}$** = $\frac{H3.3B^{K36R.}}{H3.3B^{K36R}} ; \frac{H3.3A^{2x1},\ His\Delta,\ twiGal4}{H3.3A^{2x1},\ His\Delta,\ UAS{:}2xYFP} ; \frac{12xH3.2^{K36R}}{+}$

♀ $\frac{+}{+} ; \frac{H3.3A^{2x1},\ His\Delta,\ twiGal4}{CyO}$  X

♂ $\frac{+}{} ; \frac{H3.3A^{2x1},\ His\Delta,\ UAS{:}2xYFP}{CyO} ; \frac{12xH3^{HWT}/H4^{HWT}}{12xH3^{HWT}/H4^{HWT}}$ or $\frac{12xH3.2^{K36R}}{TM6B}$

**$H3.3^{Ctrl}H4^{K16R}$** = $\frac{+}{+} ; \frac{H3.3A^{2x1},\ His\Delta,\ twiGal4}{H3.3A^{2x1},\ His\Delta,\ UAS{:}2xYFP} ; \frac{12xH4^{K16R}}{+}$

Figure S5

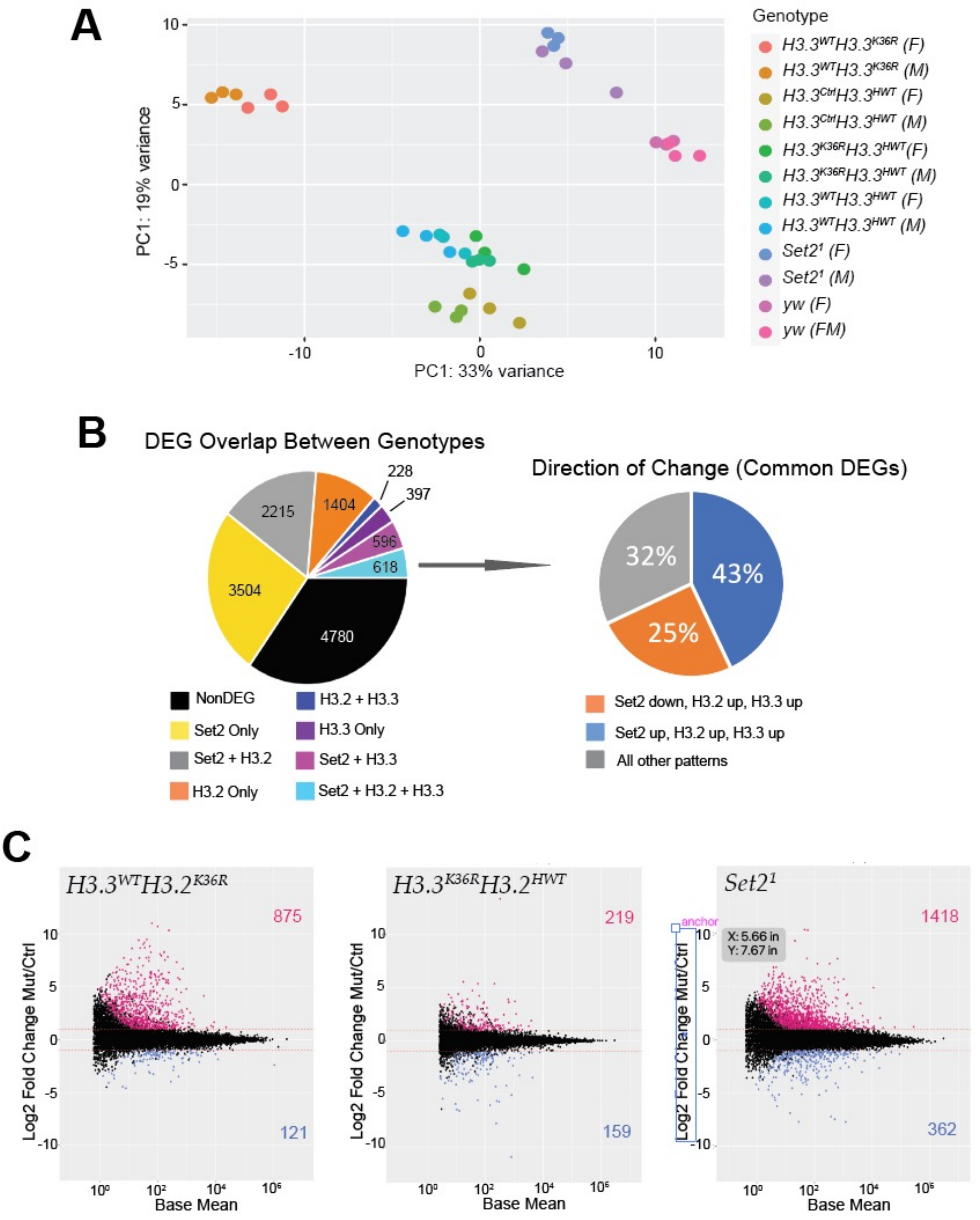

A
PC1: 19% variance
PC1: 33% variance
Genotype
H3.3WTH3.3K36R (F)
H3.3WTH3.3K36R (M)
H3.3CtrlH3.3HWT (F)
H3.3CtrlH3.3HWT (M)
H3.3K36RH3.3HWT(F)
H3.3K36RH3.3HWT (M)
H3.3WTH3.3HWT (F)
H3.3WTH3.3HWT (M)
Set2¹ (F)
Set2¹ (M)
yw (F)
yw (FM)
B
DEG Overlap Between Genotypes
228
397
2215
1404
596
618
3504
4780
Direction of Change (Common DEGs)
32%
43%
25%
NonDEG
Set2 Only
Set2 + H3.2
H3.2 Only
H3.2 + H3.3
H3.3 Only
Set2 + H3.3
Set2 + H3.2 + H3.3
Set2 down, H3.2 up, H3.3 up
Set2 up, H3.2 up, H3.3 up
All other patterns
C
H3.3WTH3.2K36R
875
121
H3.3K36RH3.2HWT
219
159
Set2¹
1418
362
anchor
X: 5.66 in
Y: 7.67 in
Log2 Fold Change Mut/Ctrl
Base Mean


Figure S6

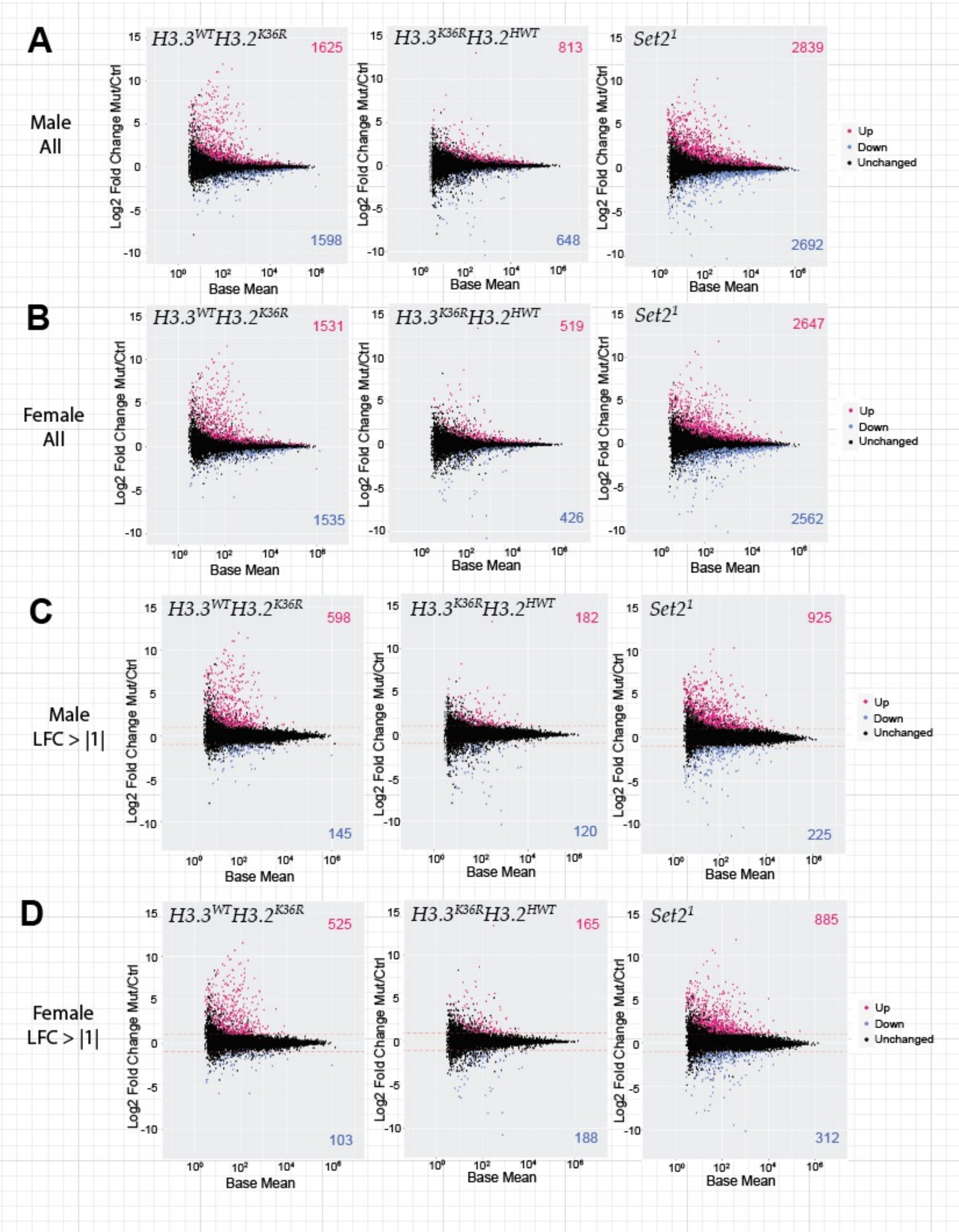
A
Male
All
H3.3^WT H3.2^K36R
1625
1598
H3.3^K36R H3.2^HWT
813
648
Set2^1
2839
2692
Up
Down
Unchanged
Log2 Fold Change Mut/Ctrl
Base Mean
B
Female
All
1531
1535
519
426
2647
2562
C
Male
LFC > |1|
598
145
182
120
925
225
D
Female
LFC > |1|
525
103
165
188
885
312

Figure S7

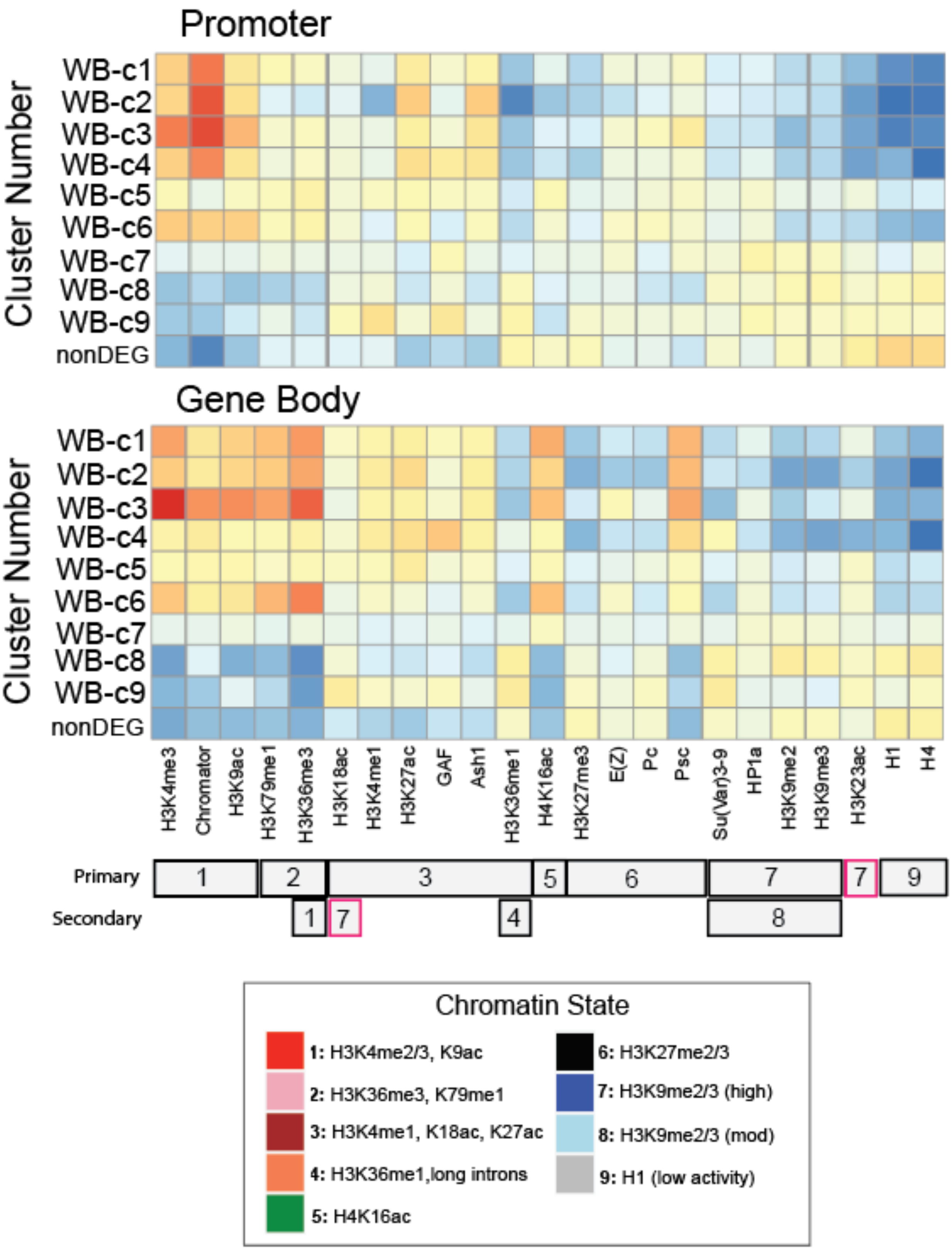
Promoter
Cluster Number
WB-c1
WB-c2
WB-c3
WB-c4
WB-c5
WB-c6
WB-c7
WB-c8
WB-c9
nonDEG
Gene Body
Cluster Number
WB-c1
WB-c2
WB-c3
WB-c4
WB-c5
WB-c6
WB-c7
WB-c8
WB-c9
nonDEG
H3K4me3
Chromator
H3K9ac
H3K79me1
H3K36me3
H3K18ac
H3K4me1
H3K27ac
GAF
Ash1
H3K36me1
H4K16ac
H3K27me3
E(Z)
Pc
Psc
Su(Var)3-9
HP1a
H3K9me2
H3K9me3
H3K23ac
H1
H4
Primary
1
2
3
5
6
7
7
9
Secondary
1
7
4
8
Chromatin State
1: H3K4me2/3, K9ac
2: H3K36me3, K79me1
3: H3K4me1, K18ac, K27ac
4: H3K36me1,long introns
5: H4K16ac
6: H3K27me2/3
7: H3K9me2/3 (high)
8: H3K9me2/3 (mod)
9: H1 (low activity)

Figure S8

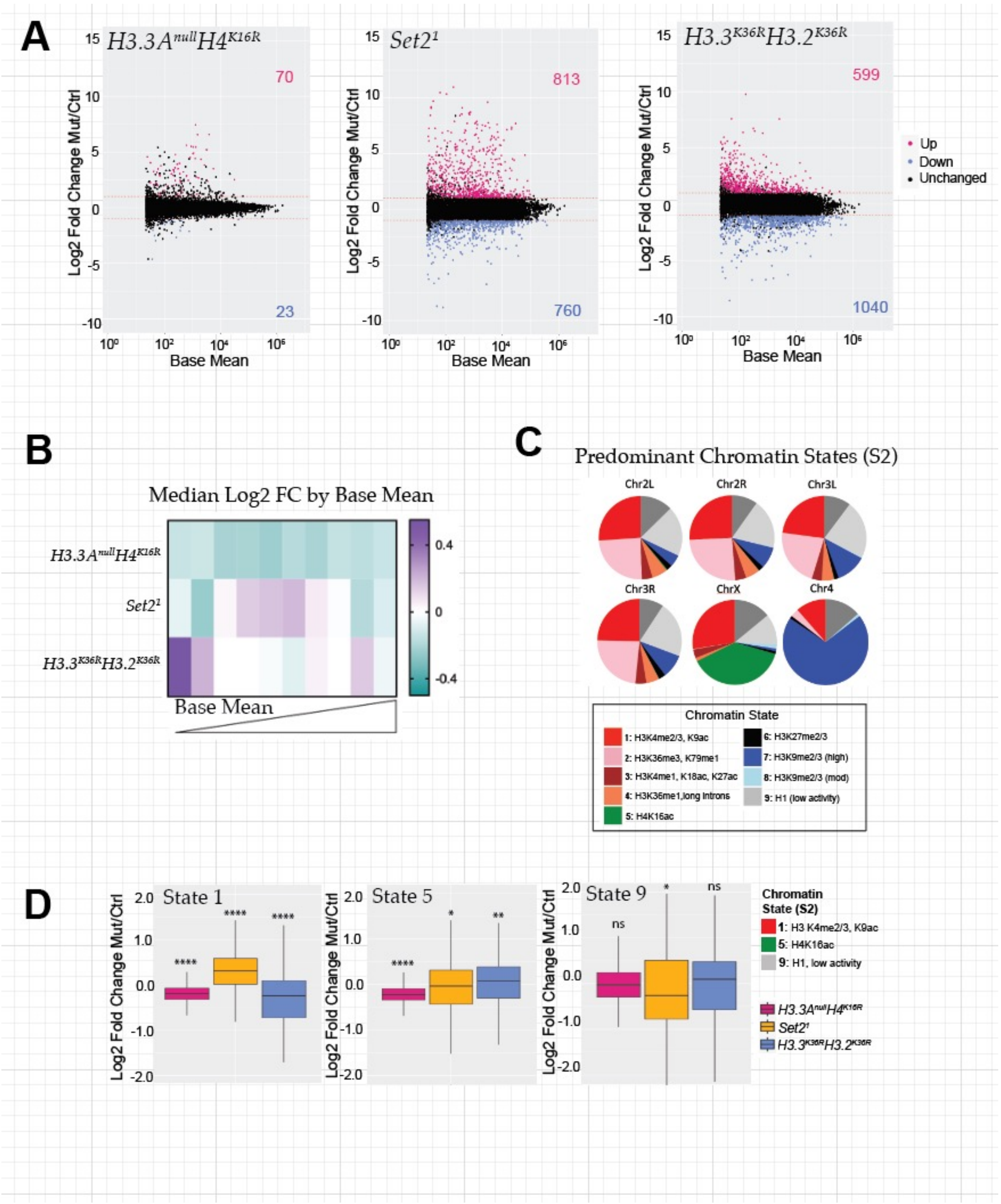
A
H3.3A^null H4^K16R
Set2^1
H3.3^K36R H3.2^K36R
Log2 Fold Change Mut/Ctrl
Base Mean
70
23
813
760
599
1040
Up
Down
Unchanged
B
Median Log2 FC by Base Mean
Base Mean
C
Predominant Chromatin States (S2)
Chr2L
Chr2R
Chr3L
Chr3R
ChrX
Chr4
Chromatin State
1: H3K4me2/3, K9ac
2: H3K36me3, K79me1
3: H3K4me1, K18ac, K27ac
4: H3K36me1,long introns
5: H4K16ac
6: H3K27me2/3
7: H3K9me2/3 (high)
8: H3K9me2/3 (mod)
9: H1 (low activity)
D
State 1
State 5
State 9
Chromatin State (S2)
1: H3 K4me2/3, K9ac
5: H4K16ac
9: H1, low activity

Figure S9

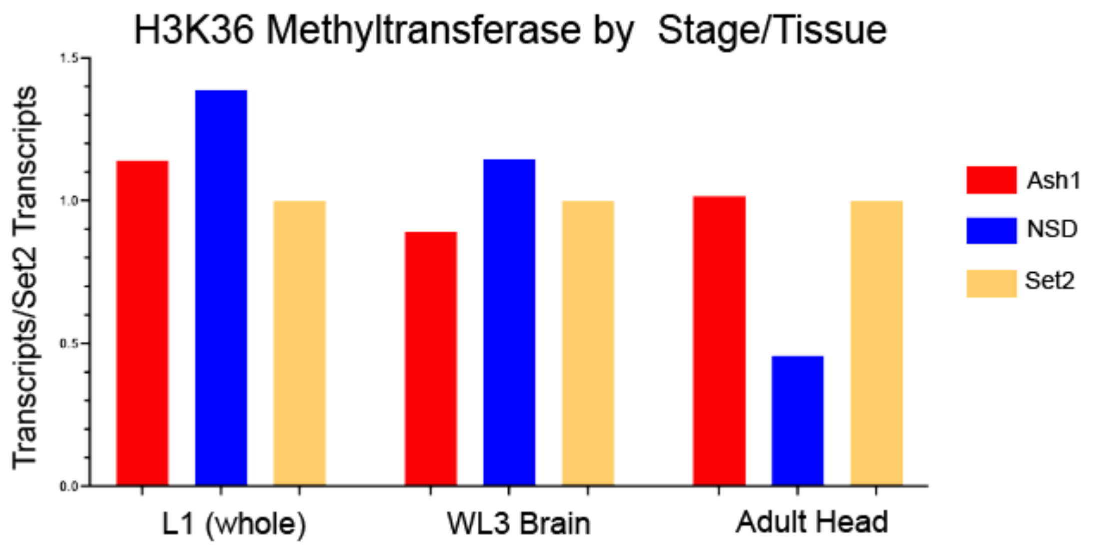
H3K36 Methyltransferase by Stage/Tissue
Transcripts/Set2 Transcripts
1.5
1.0
0.5
0.0
L1 (whole)
WL3 Brain
Adult Head
Ash1
NSD
Set2

Figure S10

#### Experimental and Control Genotype Pairs for Genomic Experiments

| Genotype | Control | Relevant Figures |
|---|---|---|
| ***$Set2^{1}$*** | ***yw*** | 2,4,5,6,7 |
| ***$H3.3^{WT}3.2^{K36R}$*** | ***$H3.3^{WT}H3.2^{HWT}$*** | 2,4,5,6 |
| ***$H3.3^{K36R}H3.2^{HWT}$*** | ***$H3.3^{Ctrl}H3.2^{HWT}$ = $H3.3^{Ctrl}H4^{HWT}$*** | 2,4,5,6 |
| ***$H3.3^{K36R}H3.2^{K36R}$*** | ***$H3.3^{Ctrl}H3.2^{HWT}$ = $H3.3^{Ctrl}H4^{HWT}$*** | 7 |
| ***$H3.3^{Ctrl}H4^{K16R}$*** | ***$H3.3^{Ctrl}H3.2^{HWT}$ = $H3.3^{Ctrl}H4^{HWT}$*** | 7 |
| ***$H3.3^{K36R}$*** | ***$H3.3^{Ctrl}$*** | 3, S2 |

Table S1

| Modification/Protein/Mutant/ Depletion | Data Type | GEO Accession Number | Cell Type/ Tissue | Figure(s) |
|---|---|---|---|---|
| H3.3$^{K36R}$ | RNA-Seq | GSE244389 | Adult Head | 3 |
| H3K36me1 | ChIP-Seq | GSE20782 | BG3 | 5D, S9 |
| H3K36me2 | ChIP-Seq | GSE51992 | BG3 | 5D, S9 |
| H3K36me3 | ChIP-Seq | GSE20783 | BG3 | 5D, S9 |
| H4K16ac | ChIP-Seq | GSE20798 | S2 | 5E, 7H |
| MOF | ChIP-Seq | GSE27806 | S2 | 5E, 7H |
| Clamp | ChIP-Seq | GSE119708 | S2 | 5E, 7H |
| MSL2 | ChIP-Seq | GSE119708 | S2 | 5E, 7H |
| MSL1 | ChIP-Seq | GSE32762 | S2 | 5E, 7H |
| MSL3 | ChIP-Seq | GSE130112 | S2 | 5E, 7H |
| Jasper | ChIP-Seq | GSE130112 | S2 | 5E, 7H |
| Jil-1 | ChIP-Seq | GSE20827 | S2 | 5E, 7H |
| H3K4me3 | ChIP-Seq | GSE20839 | BG3 | S9 |
| Chromator | ChIP-Seq | GSE20761 | BG3 | S9, 6B. 6C, 6D |
| H3K9ac | ChIP-Seq | GSE32830 | BG3 | S9 |
| H3K79me1 | ChIP-Seq | GSE32736 | BG3 | S9 |
| H3K18ac | ChIP-Seq | GSE20774 | BG3 | S9 |
| H3K4me1 | ChIP-Seq | GSE23468 | BG3 | S9 |
| H3K27ac | ChIP-Seq | GSE20778 | BG3 | S9 |
| GAF | ChIP-Seq | GSE23466 | BG3 | S9, 6B |
| Ash1 | ChIP-Seq | GSE32748 | BG3 | S9 |
| H4K16ac | ChIP-Seq | GSE20795 | BG3 | S9 |
| H3K27me3 | ChIP-Seq | GSE32791 | BG3 | S9 |
| E(Z) | ChIP-Seq | GSE27729 | BG3 | S9 |
| Pc | ChIP-Seq | GSE20803 | BG3 | S9 |
| Psc | ChIP-Seq | GSE25370 | BG3 | S9 |
| Su(Var)3-9 | ChIP-Seq | GSE20834 | BG3 | S9 |
| HP1a | ChIP-Seq | GSE23481 | BG3 | S9 |
| H3K9me2 | ChIP-Seq | GSE20791 | BG3 | S9 |
| H3K9me3 | ChIP-Seq | GSE20793 | BG3 | S9 |
| H3K23ac | ChIP-Seq | GSE20776 | BG3 | S9 |
| H1 | ChIP-Seq | GSE32767 | BG3 | S9 |
| H4 | ChIP-Seq | GSE32770 | BG3 | S9 |
| CP190 | ChIP-Seq | GSE20814 | BG3 | 6B, 6C,6D |
| BEAF-32 | ChIP-Seq | GSE32775 | BG3 | 6B, 6C,6D |
| SuHw | ChIP-Seq | GSE32810 | BG3 | 6B,6D |
| dwg | ChIP-Seq | GSE25373 | BG3 | 6B |
| CTCF | ChIP-Seq | GSE32749 | BG3 | 6B |
| Mod(mdg4) | ChIP-Seq | GSE20802 | BG3 | 6B |
| BEAF-32 RNAi | RNA-Seq | GSE147059 | BG3 | 6E, 6F |
| BEAF-32/DREF RNAi | RNA-Seq | GSE147059 | BG3 | 6E. 6F |
| CP190/Chromator RNAi | RNA-Seq | GSE147059 | BG3 | 6E, 6F |
| H3K36me1 | ChIP-Seq | GSE20782 | S2 | 7G |
| H3K36me3 | ChIP-Seq | GSE20783 | S2 | 7G |

Table S2
